# Emergence and evolution of the ERM proteins and merlin in metazoans

**DOI:** 10.1101/631770

**Authors:** V. Shabardina, Y. Kashima, Y. Suzuki, W. Makalowski

## Abstract

Ezrin, radixin, moesin, and merlin are the cytoskeletal proteins which functions are specific to metazoans. They participate in cell cortex rearrangements, including cell-cell contact formation, and play important role in cancer progression. Here we perform a comprehensive phylogenetic analysis of the proteins spanning 87 species. The results describe a possible mechanism of the protein family origin in the root of Metazoa, paralogs diversification in vertebrates and acquisition of novel functions, including tumor suppression. In addition, a merlin paralog, present in most of vertebrates, but lost in mammals, has been described. We also highlight the set of amino acid variations within the conserved motifs as the candidates for determining physiological differences between the ERM paralogs.

## Introduction

Ezrin, radixin, and moesin of the ERM protein family, further ERMs, are cytoskeleton proteins that mediate physical connection between intermembrane proteins and actin filaments (Bretscher, Edwards and Fehon, 2002). They also act as signaling molecules, for example, as intermediaries in Rho signaling (Ivetic and Ridley, 2004). Therefore, ERMs facilitate diverse cellular processes, ranging from cytoskeleton rearrangements to immunity (Bosanquet et al. 2014; McClatchey 2014; Ivetic & Ridley 2004; Marion et al. 2011). Dysregulation of ERMs’ activity and expression impairs normal wound healing process and contributes to the progression of different types of tumors (Bosanquet *et al.*, 2014; Clucas and Valderrama, 2014).

The activity of ERMs in the cell is regulated by the conformational switch from the inactive, dormant folding to the active, “stretched” form. The inactive state is established through the auto-inhibitory interaction between the N-terminal FERM and the C-terminal CERMAD domains. That results in masking of the binding sites for membrane proteins in the FERM domain and actin binding cite (ABS) in the CERMAD (Turunen *et al.*, 1998). Upon activation ERMs consequently exposed to their two activating factors: PIP_2_ (phosphatidylinositol 4,5-bisphosphate) that binds to the FERM domain and phosphorylation of a conserved threonine in CERMAD (Yonemura et al. 2002; Niggli & Rossy 2008). The important role during this transition belongs to the middle α-helical domain. In the dormant ERMs it forms coiled-coil structure, bringing N- and C-terminal domains together (Schliep *et al.*, 2017).

Little is known about individuality of each ERMs in, both, health and disease. The three proteins are paralogs and share high amino acid sequence similarity (~75% in human) (Funayama *et al.*, 1991; Lankes and Furthmayr, 1991). They demonstrate similar cellular localization and are often discussed as functionally redundant. However, data from several studies on knock-out mice revealed different phenotypes targeting different organs, and only ezrin’s depletion appeared to be lethal (Kikuchi *et al.*, 2002; Kitajiri *et al.*, 2004; Saotome, Curto and McClatchey, 2004; Liu *et al.*, 2015). Special interest in ERMs’ role in cancer stimulated multiple studies. The proteins’ dysregulation can lead to disruption of cell-cell contacts, enhanced cell migration and invasion, and higher cancer cells survival (Clucas and Valderrama, 2014). Some studies showed that ezrin, radixin and moesin may exploit different cellular mechanisms in tumors. (Pujuguet *et al.*, 2003; Kobayashi *et al.*, 2004; Debnath and Brugge, 2005; Estecha *et al.*, 2009; Chen *et al.*, 2012; Valderrama, Thevapala and Ridley, 2012).

One of the well-known tumor suppressor factors is an ERM-like protein merlin. In human, it shares 46% amino acid sequence similarity with the whole-length ezrin and 86% similarity when comparing only FERM domains (Turunen *et al.*, 1998). Mutations in merlin results in development of neurofibromatosis type 2 characterized by formation of schwannomas (Stickney *et al.*, 2004; Curto *et al.*, 2007). Tumor suppression activity of merlin is linked to the blue-box region in its FERM domain, conserved serine Ser518 and last 40 residues in the C-terminus (Lallemand, Saint-Amaux and Giovannini, 2009; Cooper and Giancotti, 2014). Two acting binding sites in merlin are located in the FERM domain, while typical to ERMs C-terminal ABS is absent (Roy, Martin and Mangeat, 1997; Brault *et al.*, 2001).

With the more research being done, it is getting clear that ezrin, radixin, and moesin can invoke different physiological effects in different tissue types, especially in cancer (Clucas and Valderrama, 2014). However, their highly conserved sequence and tertiary structure make it not a trivial task to distinguish their functions in vivo. Phylogenetic approach can be an effective tool in resolving the problem, as it enables precise paralogs characterization by tracing evolutionary history of the binding sites and conserved amino acid motifs. So far only few phylogenies of ERMs and merlin have been described in the literature. As a rule, they feature limited taxonomy representation or are included in the studies as an accessory and brief part of the discussions (Turunen *et al.*, 1998; Golovnina *et al.*, 2005; Phang *et al.*, 2016). Thus, although these phylogenies recover some interesting patterns, they do not provide the full understanding of ERMs and merlin evolution. The studies agree on the fact that the proteins are highly conserved within the metazoan clade, especially in vertebrates. Moreover, the appearance of the ERM proteins and merlin in the tree of Life seems to coincide with the origin of multicellularity in animals (Bretscher, Edwards and Fehon, 2002; Omelyanchuk *et al.*, 2009; Nambiar, McConnell and Tyska, 2010; Sebé-Pedrós *et al.*, 2013). This view is supported by the recent discovery of ERM-like proteins in Choanoflagellata and Filasterea, the closets unicellular relatives of metazoans (Fairclough *et al.*, 2013; Suga *et al.*, 2013).

The position of merlin relative to the ERM family differs in the literature, and some studies exclude merlin from the discussion of the ERM family altogether. Nevertheless, since these proteins share evolutionary history and structural characteristics, it is reasonable to unite them in one group. In this work we conduct the first comprehensive phylogenetic analysis for the ERM family and merlin, that includes data from all sequenced by the time metazoan orders. The results describe the ERM and merlin sequence conservation and paralog number diversity within the clade of Metazoa. We suggest that the increased organism complexity led to diversification of the protein paralogs in vertebrates. Moreover, we highlight the importance of phylogenetic studies of paralogs, in general, in application to experimental biology, especially in disease-related research.

## Results

### Full length ERM/merlin-like proteins appeared within Metazoa-Filasterea-Choanoflagellata group

Search for the ERM and merlin homologs throughout all eukaryotic clades resulted in the selection of 285 protein sequences spanning 87 species, including metazoan and unicellular organisms. ERM-like proteins are also present in choanoflagellates *(Salpingoeca rosetta* and *Monosiga brevicollis)* and filastereans *(Capsaspora owczarzaki).* A sequence of 298 amino acids (Supplementary file) identified in corallochytrean *Corallochytrium limacisporum* revealed 25% sequence identity to the FERM domain of human ezrin based on the tBLASTn search against the species’ whole genome sequence. Although such similarity level does not guarantee structural alikeness, according to (Rost and Sander, 1994), the domain annotation by PFAM indicated that this polypeptide from *C. limacisporum* belongs to the class of FERM domain with the high statistical support (e-value < 10^-5^ for each of the three subdomains of FERM). Sequence-based prediction of biochemical properties revealed that the two FERM domains, human and corallochytrean, exhibit distinct features, including pI (8.75 for human ezrin FERM domain and 6.79 for the *C. limacisporum* polypeptide), amino acid content, instability index and hydropathicity. In particular, metazoan FERM domain is predicted to be more hydrophilic than its suggested corallochytrean homolog (grand average of hydropathicity (GRAVY) index is – 0.530 and −0.270, respectively) and less stable (instability index estimated to be 43.57, i.e. unstable protein, and 31.80, stable, respectively). The only binding site conserved in the corallochytrean FERM is the site for PIP_2_ interaction. No ERM-like proteins or FERM-like domains could be found in the other inspected taxa. The list of all the taxa and the corresponding proteins IDs used for the analysis can be found at the Supplementary (Table S2).

### ERM+merlin protein family is conserved throughout all the metazoan orders and three unicellular species

Sequence comparison of the selected proteins demonstrated that the domain structure and most of the known binding sites are conserved throughout the whole metazoan clade, although there is some length variation. The amino acid motifs conservation analysis (Fig. 1) revealed that the homologs of such early branching animals as *Trixhoplax adhaerens* (Placozoa) and *Amphimedon queenslandica* (Porifera) demonstrate similar pattern to the mammalian proteins with a high statistical significance. The FERM and CERMAD domains are characteristically well preserved, even in the proteins from the unicellular organisms, while α-helical middle domain is the least conserved ERM domain as has been previously described (Phang *et al.*, 2016). The N-terminal part of the FERM shows some length variation throughout different taxa by including short, non-conserved amino acid stretches. More striking length variation can be observed within the region separating the α-helical domain and the CERMAD. It is short for proteins from vertebrate animals, but is increased in all other taxa, the longest is in the protein from *C. owczarzaki.* The sequence of this region is poorly preserved between the taxa. The proteins from the species of flat worms, *Inoshia linei* and *C. owczarzaki* are the most divergent.

**Figure 1.**
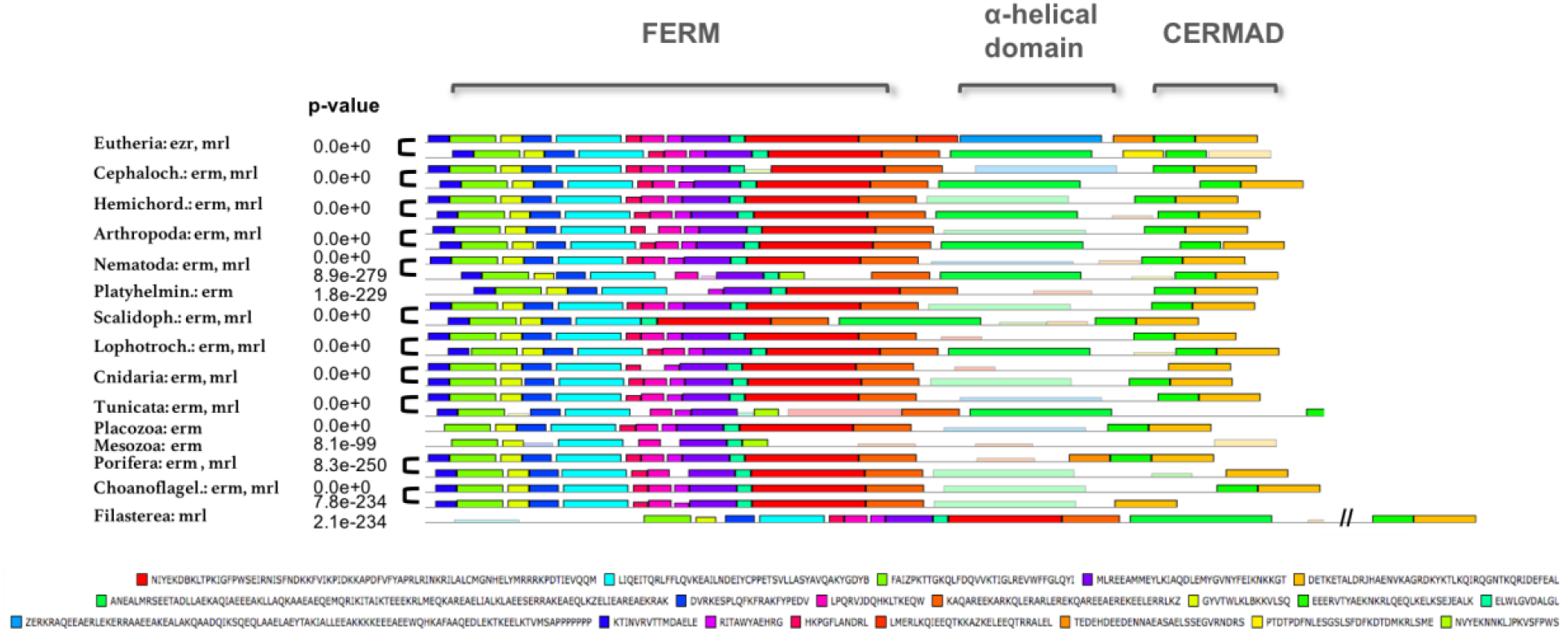
MEME conserved amino acid motif analysis. Motifs in pale color were found by the scanning algorithm based on the de-novo motif identification (bright color). Note the reduced length of the region separating the alpha-helical and CERMAD domains in eutherian proteins.

The binding sites for EBP50, ICAM-2, NHERF2 binding partners of the ERMs, for PIP_2_, ABS, and intramolecular interaction sites (Bretscher et al. 2002; McClatchey 2014; Ivetic & Ridley 2004; Marion et al. 2011; Li et al. 2015) can be identified in the most of the sequences (Supplementary, Table S2). This signifies that the protein family is responsible for the basic cellular functions and some of the interactions might have been established in the early metazoan history. The most conserved regions in the ERM proteins are PIP_2_ binding sites in F1 and F3 subdomains of the FERM domain and the C-terminal ABS (specifically, KYKTL motif). The most differing proteins come from the early branching metazoans *Molgula tectiformis* (Tunicata), *A. queenslandica* (PoriferaJ and endo-and exo-parasites (*I. linei,* flat worms, and blood sucking leech). Interestingly PIP_2_ binding motif within F1 subdomain is not preserved only in *M. tectiformis.* Small variation in KYKTL motif can be seen in the proteins from Echinodermata and shark *Callorhinchus milii.*

Based on the sequence, merlin proteins can be classified into three groups (Fig. 2): 1) non-vertebrate proteins; 2) vertebrate merlin1, absent in most of Eutheria 3) all-vertebrate merlin2 (we arbitrary assign here merlin1 and merlin2 names to the merlin paralogs). The group 2 comprises merlin1 proteins coded by the paralogous gene that has not been described before (the list of the proteins is in the Supplementary, Table S2). It is characterized by a unique insertion of 68 amino acids in tetrapods and 15 amino acids PPYxPHSNRNSAYMx in bonny fishes in C-terminal domain and lacks tumor-suppression region characterized in human merlin2 between residues 532-579. It does have merlin specific blue-box region and the conserved Ser518. Interestingly, within Eutheria clade only armadillos (superorder Xenarthra) possess merlin1 gene. Invertebrate merlins, group 1, are similar to vertebrate merlin1 but lack the C-terminal domain insertion. Merlin2, group 3, is present in all vertebrate taxa and has an additional actin binding site in the F1 of FERM and the tumor-suppression amino acid stretch in its CERMAD domain (residues 532-579 in human). The blue-box region can also be identified in the two proteins from the unicellular species: XP_004364665.2 in *C. owczarzaki* and XP_004991962.1 in *S. rosetta.*

**Figure 2.**
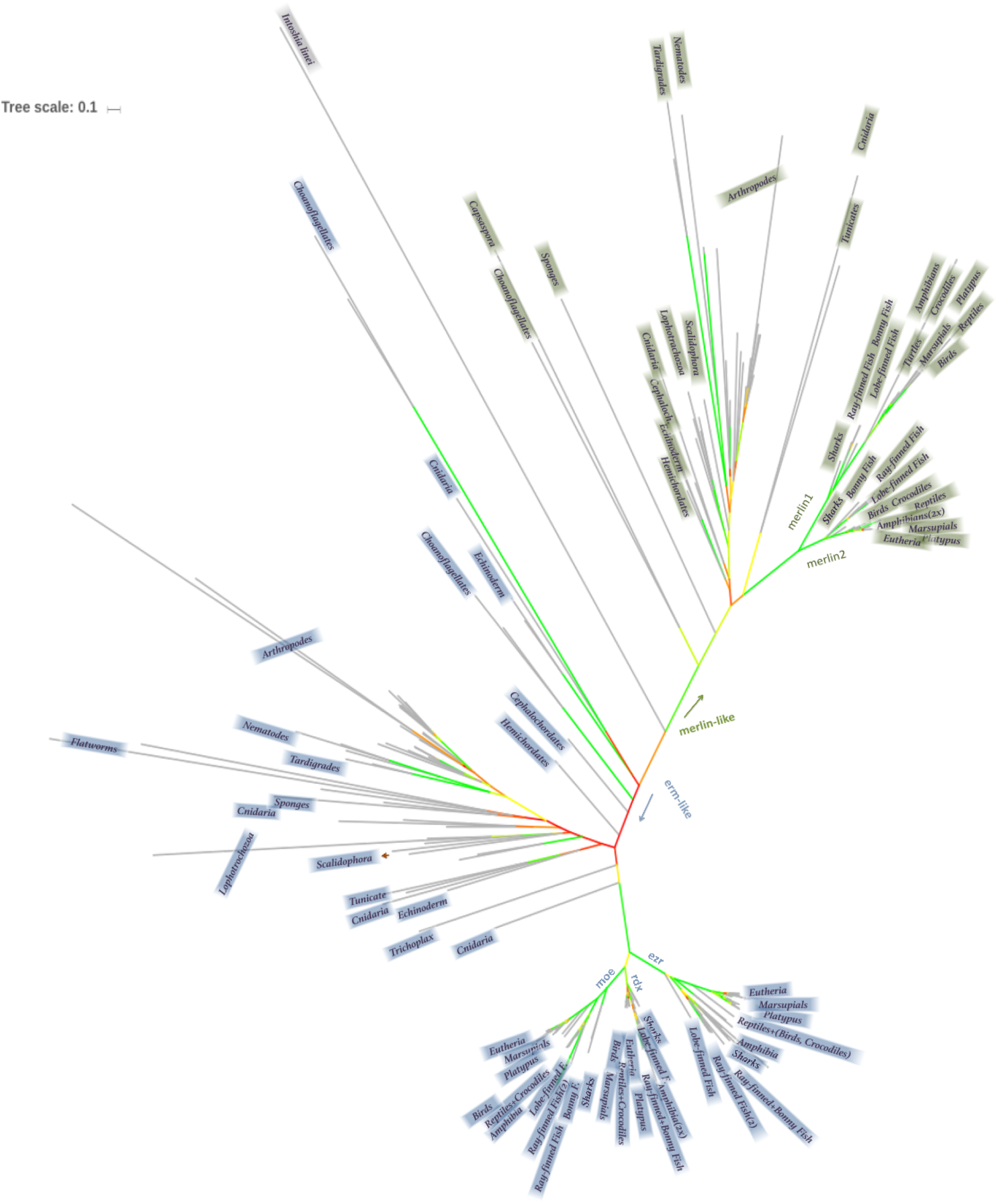
Phylogenetic ML tree: ERM+merlin family. The ERM-like part of the tree is shaded in blue, the merlin-like part is shaded in green. Note the emergence of the ezrin, radixin, moesin paralogs in vertebrates; and divergence of vertebrate merlins on merlin1 and merlin2 paralogs. Some proteins from the unicellular choanoflagellates species and a proteins from *Capsaspora* relate more to merlin-like proteins, some – to ERM-like proteins. Color scheme for bootstrap values: green – 70-100%, yellow – 50-70%, red – below 50%. Refer to the tree file in Newick format to see branch lengths and bootstrap values (Supplementary file).

### Phylogenetic tree of the ERM+merlin family

The reconstructed ML tree resulted in high phylogenetic resolution among vertebrates, while the branching for most of the other taxa have low statistical support (Fig. 2). The alternative methods of phylogenetic reconstruction, including neighbor-joining algorithm, parsimony method, and Bayesian inference, could not improve the resolution (data not shown). Two former trees revealed almost identical branching and statistical confidence. The Bayesian reconstruction could not achieve conversion after 3000000 generations. The run was terminated and the consensus tree was built anyway. It featured clustering of vertebrate proteins supported by high bootstrap values, but unresolved branching for the invertebrate sequences. One of the reasons can be an unequal representation of the taxa due to the lack of sequencing data for the invertebrate clades. Another complication for the analysis could be the high divergence of the amino acid sequences between evolutionary distant lineages. The ML tree is used for the further discussion.

The most “eccentric” sequence in the reconstructed phylogeny is the one of the hypothetical protein from *I. linei,* that, although features all three ERM domains, has the highest substitution rate and does not cluster with any other groups. Consequently, it cannot be defined as ERM-like or as merlin-like protein. This is not surprising, as *I. linei,* a representative of orthonectids, is a parasitic animal, and its hermaphrodite nature, fast reproductive cycles and high level of inbreeding can be the reason that makes its genome distant from the genomes of other metazoans (Lu *et al.*, 2017).

The protein from *I. linei* was arbitrary chosen to separate the tree into two major clusters: ERM-like and merlin-like groups. Therefore, any “hypothetical” or “unknown” proteins could be annotated whether as ERM-like or as merlin-like, based on their position relative to *I. linei*’s protein. Besides improving annotation of such proteins, some false annotations deposited in the public protein databases were corrected (Supplementary, Table S2).

Interesting observation can be made regarding the proteins from the unicellular species. The *C. owczarzaki’s* protein XP_004364665.2 and S. rosetta’s protein XP_004991962.1 cluster as merlins; and XP_004997754.1 from *S. rosetta* and XP_001746613.1 from *M. brevicollis* cluster as ERM-like. However, the bootstrap support is not high enough to confidently assign these annotations. At the same time, the proteins XP_004994097.1 from *S. rosetta* and XP_001743289.1 from *M. brevicollis* rather cluster together with the invertebrate ERM-like group. That provides an insight into the origin of the ERM+merlin family and suggests that merlin and ERMs diverged from the common ancestral protein before emergence of Metazoa.

Cnidaria, Placozoa, Echinodermata, Scalidofora, Lophotrochozoa, Porifera, Hemichordata, Cephalochordata, and Mesozoa taxa branching could not be defined by this analysis with statistical confidence. ERM-like proteins from the clade Tunicata are the most related to the vertebrate ERMs (bootstrap support 83%), but the tunicate’s merlins do not form clear clustering. The three clearly distinguishable groups, beside vertebrates, are Nematoda (99%) and Tardigrada (100%), and Arthropoda (76%) that includes Insecta (95%). ERM-like proteins from nematodes and tardigrades cluster together with the bootstrap support of 48%. The phylogenetic position of tardigrades is so far unclear and developmental stages and genetics of these animals share features of both, arthropods and round worms (Gabriel *et al.*, 2007; Yoshida *et al.*, 2017).

Ezrin divergence from the group radixin-moesin in Vertebrata is supported with 100% bootstrap value; radixin and moesin diverge in the separated clusters with the bootstrap of 66% and 100%, respectively. Radixin seems to be the slowest evolving protein in the family judging from the branch lengths. Coelacanthimorpha *(Latimeria chalumnae),* Teleostei (bonny fishes), Holostei *(Lepisosteus oculatus),* Chondrichthyes (cartilaginous fishes) and Amphibia are clearly separated from each other and the closely related group of Prototheria-Metatheria-Theria-Sauria (mammals, reptiles, birds). This is true for each of ezrin, radixin, and moesin clusters. The pattern is similar for the vertebrate merlins, except the merlins from *Xenopus laevis* cluster within the mammals-reptiles-birds group. Each ERM protein in the bonny fishes has an additional paralog and the divergence of ezrin, radixin, and moesin into two clusters each can be distinctly seen within the bonny fishes clade. This agrees with the hypothesis of the lineage-specific whole genome duplication (WGD) events (see further).

### Positive selection in fish lineages

The long branch lengths for ezrin and merlin1 proteins in vertebrates suggest that they evolved faster than radixin, moesin and merlin paralogs2. To take a deeper look into the evolution of vertebrate ezrin and merlins we analyzed the ratio of nonsynonymous to synonymous nucleotide substitutions, omega, along ezrin and merlin trees separately from the rest of the data set. The codeml test in PAML for varying omega rates among the tree branches (branch free-ratio model) verified that the varying ratio scenario is more likely than not (p-value 5.7e-109 for ezrin and 1.5e-39 for merlin1-2). The branch free-ratio model is useful for overall estimation but not very informative due to the little statistical power, as it uses a big number of parameters for calculations. Therefore, we performed as well the branch-site model analysis in order to test for positive selection within some lineages, based on the ML tree and the branch-free ratio model estimation.

Among ezrin proteins only holostei and teleost fishes are likely to have undergone evolution under positive selection (p-value 0.005). Other lineages seem to evolve under stricter constrain. Such phenomena can be explained by the additional duplication events in fishes and, therefore, weaker selective pressure (Glasauer and Neuhauss, 2014).

In the case of merlin, all tested branches of merlin1 showed evidence for positive selection (p-value 4.6e-09). At the same time the sequence of merlin2, the paralog responsible for anti-tumor activity, seems to be highly conserved during evolution in all vertebrates.

### Paralog number diversity

All invertebrate taxa included in this analysis are characterized by the presence of zero or two ERM-like paralogs and zero or one merlin-like paralogs (Table 1, Supplementary table S1). In particular, *I. linei, T. adhaerens* and all three species of Platyleminthes have only one gene coding for an ERM-like protein and no merlins. Among Nematoda species, *Loa loa* has only ERM-like protein, no merlin. These observations highlight the trend of simplification in parasites. All other invertebrate taxa have at least one ERM-like and one merlin-like proteins. Although, some paralogs can be missing from our data due to the incomplete genome assemblies. It is interesting to note that some cnidarians and lophotrochozoans underwent a local duplication of ERM-like genes, that resulted in the paralogous genes located within few thousand nucleotides from each other *(Lingula anatina, Exaiptasia pallida).*

**Table 1.**
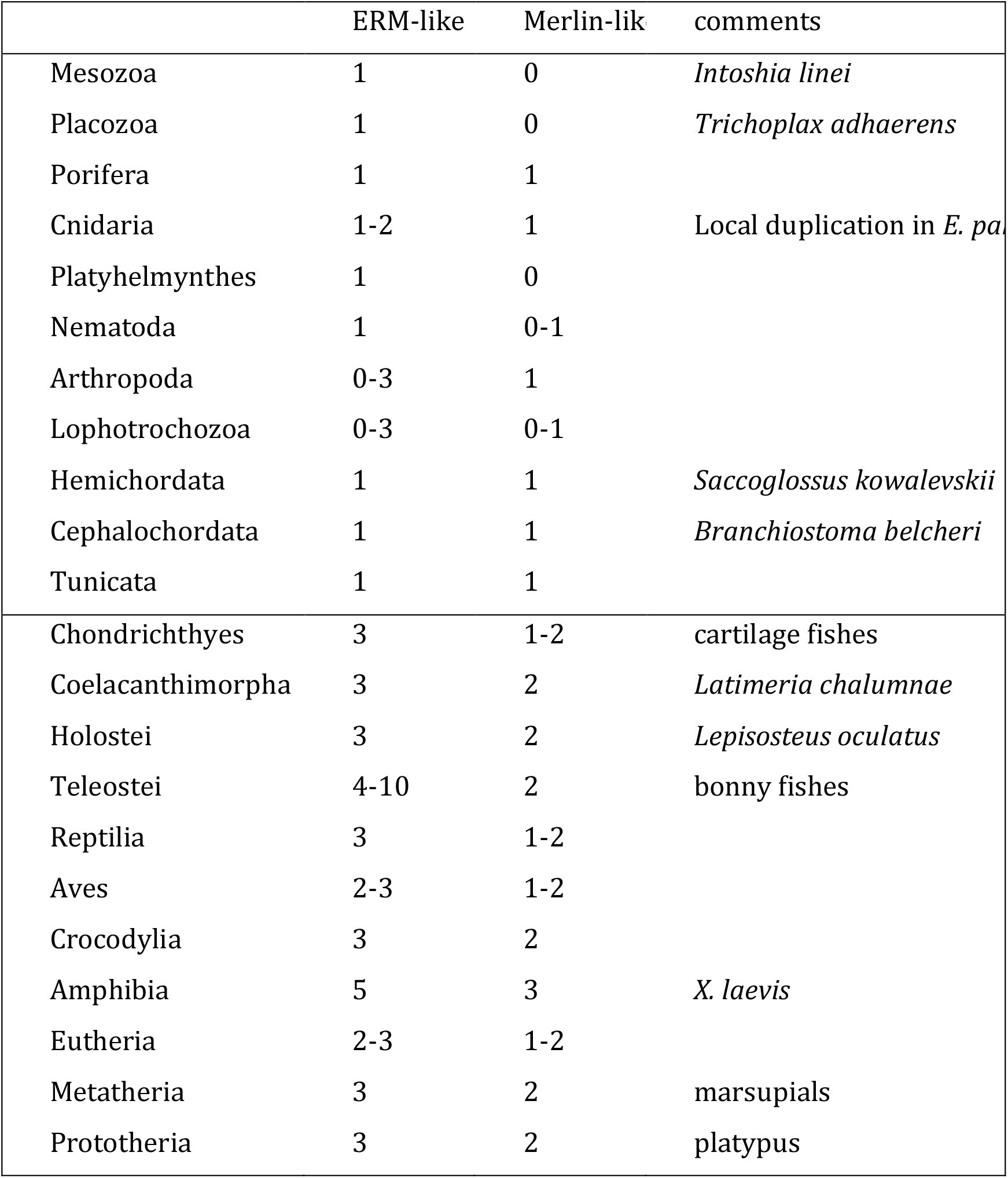
Number of paralogous genes in different metazoan lineages. For more detailed count refer to the Table S1 in the Supplementary.

Vertebrates have, as a rule, three ERM proteins (ezrin, radixin, and moesin) and two merlins. Although several very interesting exceptions can be found. As it was already mentioned, Eutheria, except armadillos, keep only merlin2. We indicated a few more cases of lineage specific paralogs lost or gain. For example, teleost fishes have four to six ERM paralogs in different combinations, and Atlantic salmon *(Salmo salar)* has ten ERM genes (two of ezrin, four radixin, and four of moesin). This observation is in accordance with the hypothesis of a lineage specific WGD in salmonids (Glasauer and Neuhauss, 2014). Further, some of Neognathae birds lost moesin gene. *X. laevis* has five ERM proteins (one ezrin, two radixins, two moesins) and three merlin paralogs, likely the result of another lineage specific WGD that took place around 40 million years ago (Van de Peer, Maere and Meyer, 2009). Elephant shrew *(Elephantus edwardii*) has two ezrin genes and one radixin. Interestingly, one of its ezrin genes is a retrogene. Besides there are at least four pseudogenes descendant from the ERM family in this species. There are few cases of ERM pseudogenes throughout mammals, but the case of elephant shrew is the most prominent.

To avoid any errors during the paralogs number estimation, we ran tBLASTn searches against any available RSA sequences and whole genome sequences for the species that show lack of any of the paralogs.

Emergence of the three ERM paralogous genes in vertebrates could be a result of the two rounds of the WGD that took place in the root of vertebrates and a consequent loss of one copy of the gene. An additional increase of the paralogs number in the teleost fishes is likely to be the result of the lineage specific WGD event. Although the possibility of a duplication event of a local character cannot be excluded.

The situation with merlin genes is different. Even the lineages that underwent the additional WGD rounds and have increased number of ERM paralogs, still keep strictly two merlin genes. However, there is a large exception within the Eutheria clade: majority of the lineages there lost merlin1 gene (Fig. 3). This fact was previously unknown or unappreciated, likely because most of the merlin studies were done on the representatives of Eutheria clade (human, mouse), thus describing only merlin2 paralog. Therefore, it is not surprising that merlin1 went unnoticed.

**Figure 3.**
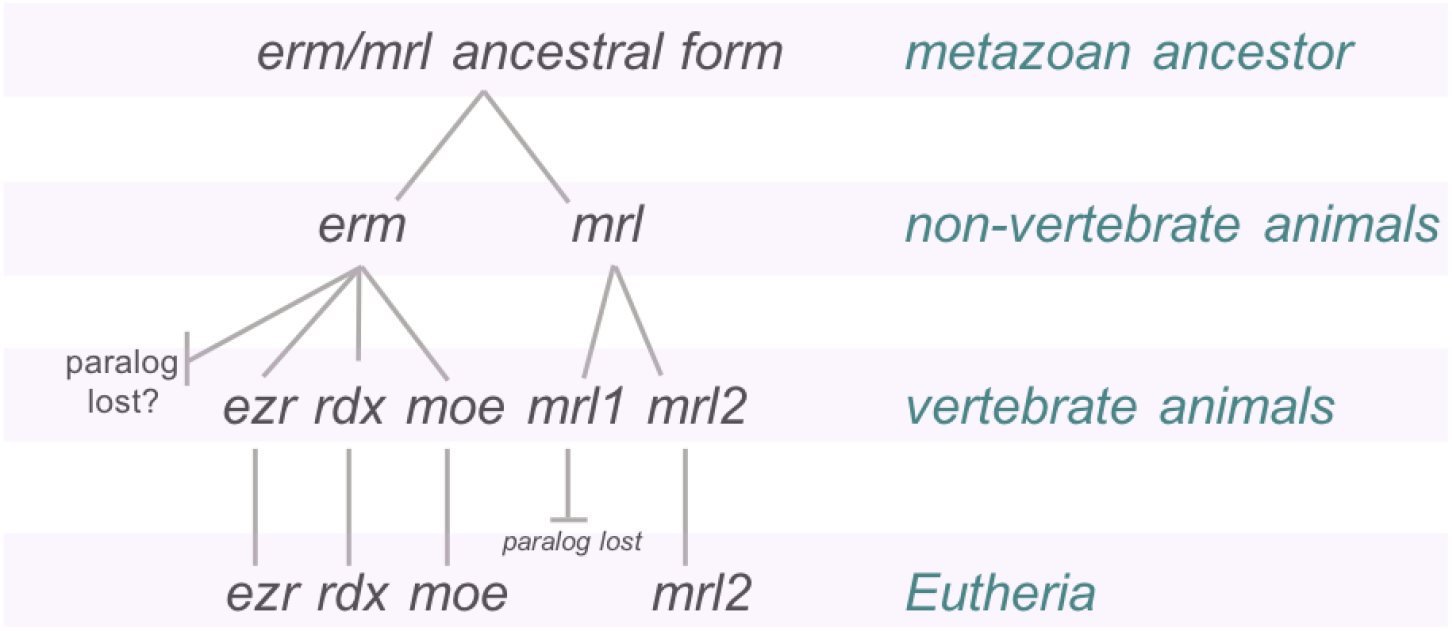
Scheme of gene duplication and paralogs lost in vertebrates. Erm – ERM-like, mrl – merlin-like, ezr – ezrin, rdx – radixin, moe – moesin.

### Syn teny rela tionships

We analyzed synteny conservation for ERM proteins at two levels: intra-synteny for ezrin, radixin, and moesin within *Homo sapiens* species, and synteny for ERMs and ERM-like proteins between metazoan lineages. Interestingly, ezrin, radixin, and moesin genes very poorly preserve synteny, neither in their closest neighborhood (~500 Gb up- and downstream), nor on the whole chromosomes (Fig. 4). Such observation suggests that the appearance of the three paralogs can be of a local character, rather than a result of the WGD event. However, in such case the persistence of the three paralogs in all the diverse vertebrate lineages would be also difficult to explain.

**Figure 4.**
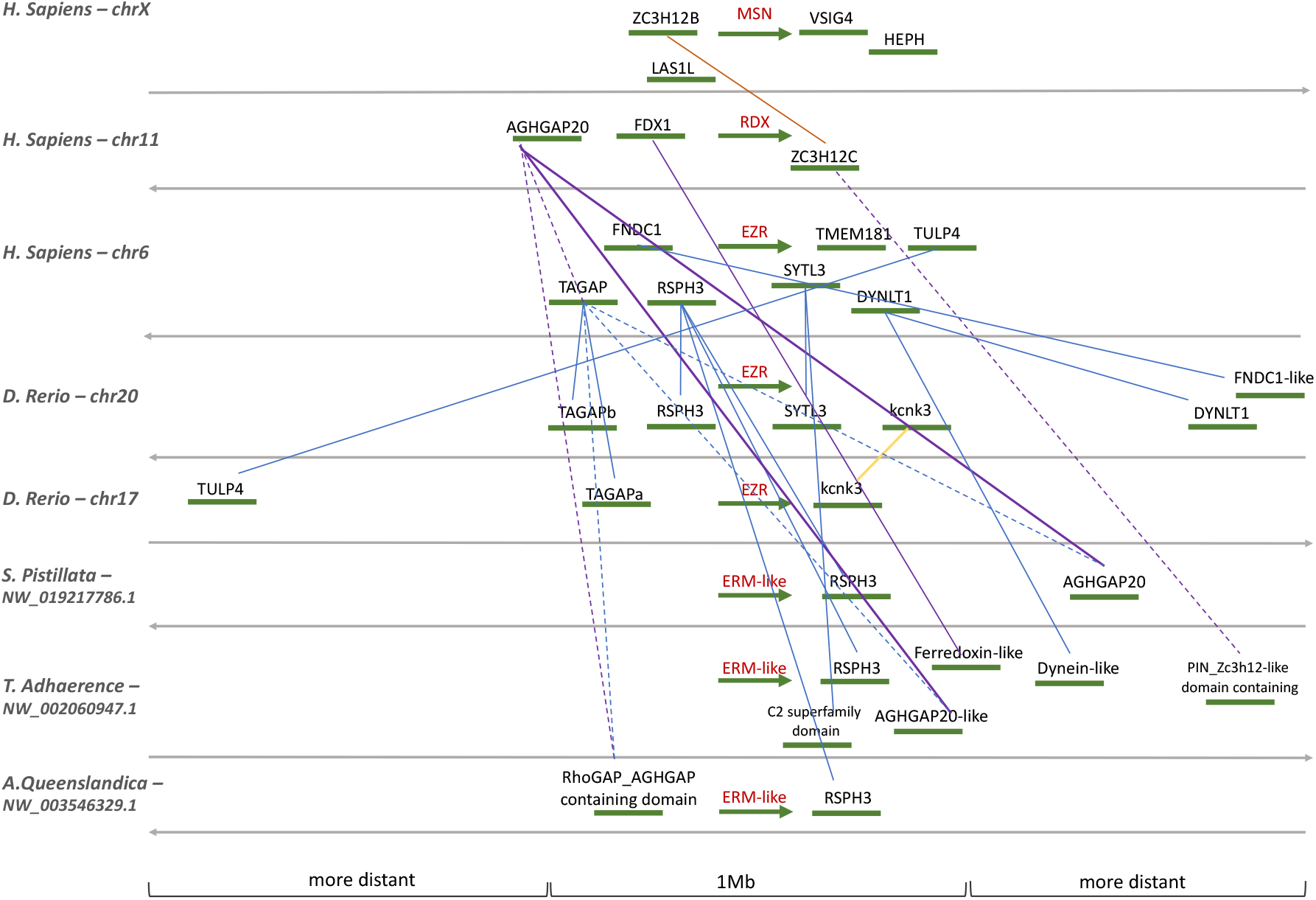
Synteny relationships for ezrin, radixin, and moesin genes in the genome of *H. sapiens* and of ezrin and ERM-like proteins in different animals. Color scheme is used for the better visualization. Solid lines connect syntenic genes, dash lines are used in case if only parts of genes (proteins domains) are syntenic.

On the other hand, the similarity can be found in genes location between mammals *(H. sapiens, Mus musculus),* fish *(Danio rerio),* cnidarians *(Stylophora pistillata), T. adhaerens,* and *A. queenslandica*. The remarkable conservation of the syntenic block can be seen between human and mouse, for example, where all seven protein coding genes surrounding ezrin in human match those in mouse (data not shown). Insects *(Drosophila melanogaster),* however, show no trace of synteny with any of the lineages, that is surprising regarding the high sequence similarity of the ERM-like protein in fruit fly and human ERMs. Such divergence could reflect the specificity of insects’ – or fruit fly, in particular – genome architecture and evolution. Perhaps, analysis of conservation of non-coding sequences, including repetitive elements, could be useful in understanding the syntenic divergence in *D. melanogaster* and character of duplication events in vertebrate lineage.

To assess the evolutionary background of the two merlin paralogs, we compared gene content around merlin1 and merlin2 genes in chicken and zebrafish. The revealed synteny suggests the higher genomic structure conservation between regions containing merlin2 genes. In particular, more syntenic genes located within 1 Mb interval and shared higher sequence identity. Synteny between the two different merlin paralogs (merlin1 and merlin2) is the least conserved, with only one syntenic gene pare present in the close proximity to the merlin genes (Tab. 2). This pattern is more striking if comparing human and zebrafish. The data present in the Synteny Database show 19 synteny pares around merlin2 gene in human and zebrafish and no similarity when comparing human merlin2 and zebrafish merlin1 (see Supplementary, Table S2).

**Table 2.** Comparison of synteny between merlin1 and merlin2 genes.

|  | Da.re mrl1-Da.re mrl2 | Da.re mrl1-Ga.gl mrl1 | Ga.gl mrl2-Da.re mrl2 |
| --- | --- | --- | --- |
| Number of<br>syntenic pares | 14 | 16 | 34 |
| Syteny within<br>1 Mb interval | 1 | 2 | 9 |
| Syntheny: same<br>chromosome | 13 | 14 | 25 |
| Sequence identi<br>estimator<sup>a</sup> | 51% | 66% | 67% |

<sup>a</sup> Sequence identity estimator shows how much of sequence identity share 50% of all syntenic pares. The detailed comparison table can be found at the Supplementary table S2.

### Unicellular ancestry of ERM+merlin family

With the assumption that the closest unicellular relatives of animals are choanoflagellates and filastereans, we assumed that their ERM-like and merlin-like proteins are the best candidates for speculating about the proteins ancestral form. We built a small phylogeny tree, including only few sequences from an eutherian representative *(H. sapiens)* and *S. rosetta, M. brevicollis, and C. owczarzaki.* Eliminating the rest of the taxa decreases the reliability of the reconstruction, but, regarding the high sequence diversity and scarcity of the data for invertebrates, this approach is the most straightforward. The tree highlights three groups: 1) merlins (Eutheria, Choanoflagellata, Filasterea), although this cluster can be separated into two subgroups: filasteran and eutarian+choanoflagellate; 2) highly specialized (or the result of genome mis-assembly) proteins of the two choanolagellate short proteins lacking most of the middle domain; 3) ERM group comprising eutherian and choanoflagellate homologs (Fig. 5).

**Figure 5.**
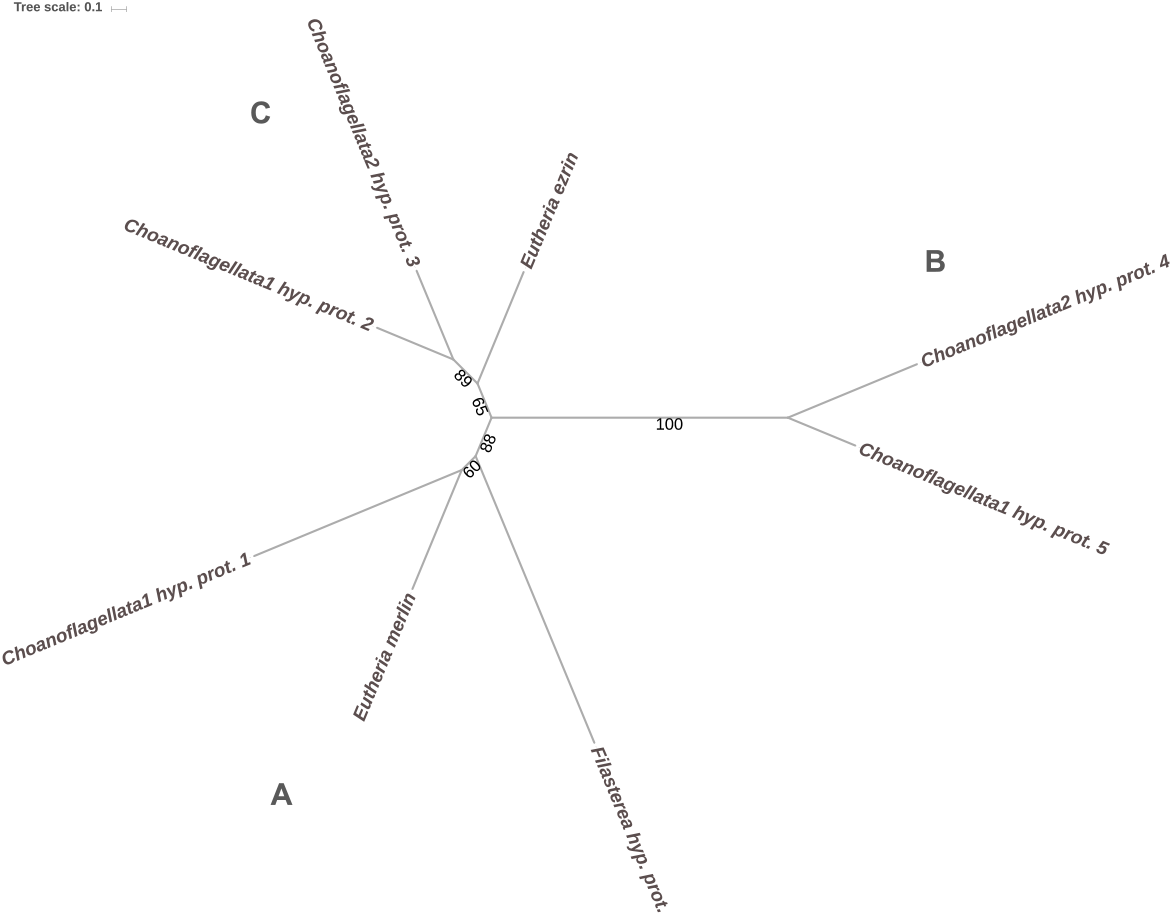
ML tree indicating three clusters of ERM/merlin-like proteins in the unicellular organisms: two species form choanoflagellates and one filasterean species. Group A unites merlin and merlin-like proteins, clade B is represented by the two proteins with a rudimental middle domain, clade C – ERM-like proteins. The tree in Newick format is in Supplementary file.

Although the tree is more a scheme than an illustration of the phylogenetic relationships, it is useful for speculating about the origin of the modern protein family. We, therefore, modelled an ancestral sequence for ERM+merlin family based on this tree (Supplementary file). It suggests the conserved domain structure and presence of the most characteristic binding sites (for PIP_2_, intramolecular interaction, ABS), but includes an additional 63 amino acid region separating alpha-helical domain and CERMAD. Computational prediction of the tertiary structure of this insertion characterized the corresponding polypeptide as an extended structure with a low probability to form α-helix (Fig. 6).

**Figure 6.**
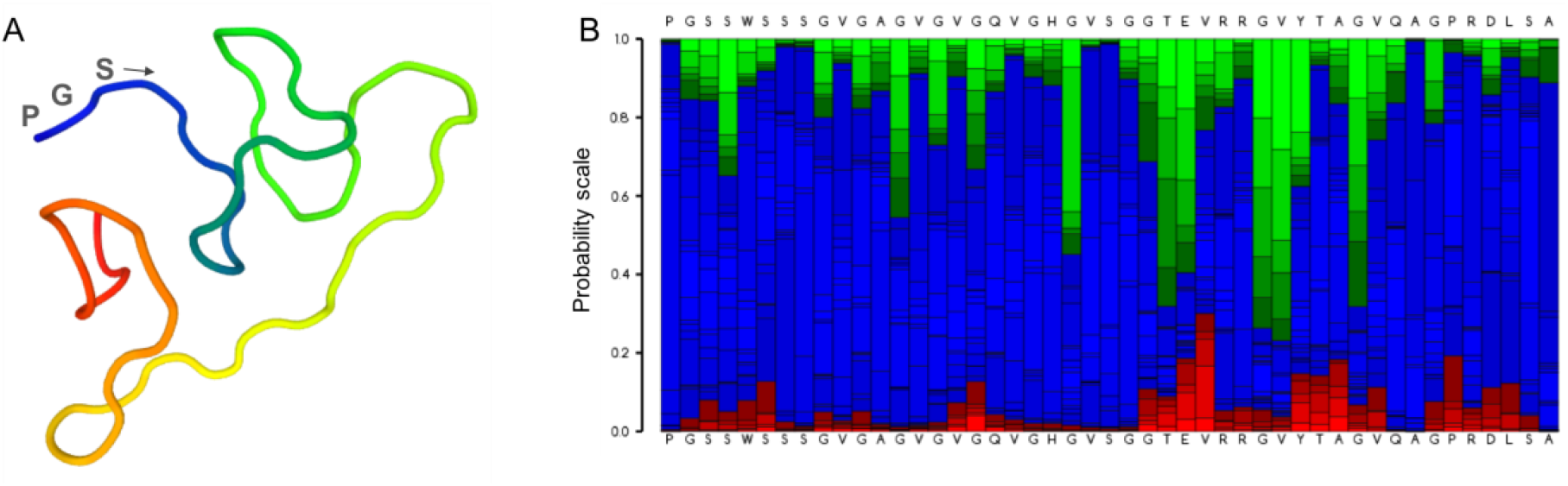
Structural modelling. A: 3D peptide structure prediction for the first 50 residues of the insertion in the reconstructed ancestral protein. Coloring is used for a better visualization. B: Probability plot for the first 47 amino acids of the insertion, each amino acid is assigned a probability to be included in a particular structure: red – α-helix, blue – random coil, green – extended structure. Higher values mean higher probability. The plot suggests that the analyzed polypeptide folds into an extended structure. Reconstruction was done in PEP-FOLD3.

## Discussion

Phylogenetic analysis is a valuable approach in protein annotation and characterization and can significantly improve genome annotations. Unfortunately, it requires more time and efforts than an automated genome annotation pipelines. However, it can and should be routinely used for proteins that are actively studied in vivo and for medical applications, if not for evolutionary studies. The presented here phylogenetic tree allows distinguishing between ERM-like and merlin-like subgroups of the protein family and between different ERM paralogs in the approach proposed more than 20 years ago by Jonathan Eisen (Eisen, 1998). As a result, we were able to improve annotations of these proteins in different species, as well correcting errors in several cases when, for example, an ezrin was erroneously called a radixin or a moesin.

Furthermore, the tree clearly demonstrates that assignment of the invertebrate proteins to “ezrin”, “radixin” or “moesin” is inconsistent, since the divergence of the three ERM paralogs happened in the root of Vertebrata. A good example are the two incorrect assignments at the NCBI protein database: XP_002160112.1 protein annotated as radixin in *Hydra vulgaris* and NP_727290.1 protein annotated as moesin in *D. melanogaster.* We suggest to restrict the names ezrin, radixin, moesin only to the vertebrate proteins, while referring to others as ERM-like or merlin-like.

Some inconsistencies also happen in the studies with vertebrate ERMs. For example, there are experimental data discussed for chicken moesin (Winckler *et al.*, 1994; Li and Crouch, 2000), even though the authors did not find enough evidence to confidently claim moesin’s expression in the samples. Indeed, such experiments are questionable, as our phylogenetic analysis indicates that chicken *(Gallus gallus)* lost moesin gene and has only ezrin and radixin. Such an example shows the importance of incorporating bioinformatics milieu in the wet-lab studies, especially for the proteins from the less studied, not model organisms, to avoid confusion and data discrepancy.

To get an insight on the evolution of the ERM+merlin protein family, we collected protein sequences from 84 species of Metazoa and 3 unicellular species from the clades Choanoflagellata and Filasterea. We could also identify a FERM-like domain with the conserved PIP_2_ binding site in a species from Corallochytrea clade, likely the first lineage with FERM domain in the tree of Life (Fig. 7A). This finding is in agreement with the study of the domain gain and lost in different taxa, that estimated that the FERM’s origin took place in Holozoa (Grau-Bové *et al.*, 2017). This is also consistent with the fact that FERM domain is the most conserved part of the proteins in the family. It is possible, that this corallochytrean FERM domain is a full-length protein, as computational prediction by ExPASy ProtParam algorithm estimated that it likely folds into a stable structure. We suggest that this FERM-like protein is a membrane binding protein involved in communication of the cell and extracellular environment. It can be similar to a predecessor of the ERM+merlin family. Domain shuffling and/or shifts within the open reading frame of the predecessor gene could lead to the origin of a longer protein with an acting binding capability of its newly acquired C-terminal part, i.e. with a scaffolding function similar to that of the modern ERMs. Such mechanisms, for example, were discussed in the studies about emergence of animal multicellularity (Shalchian-Tabrizi *et al.*, 2008; Richter and King, 2013).

**Figure 7.**
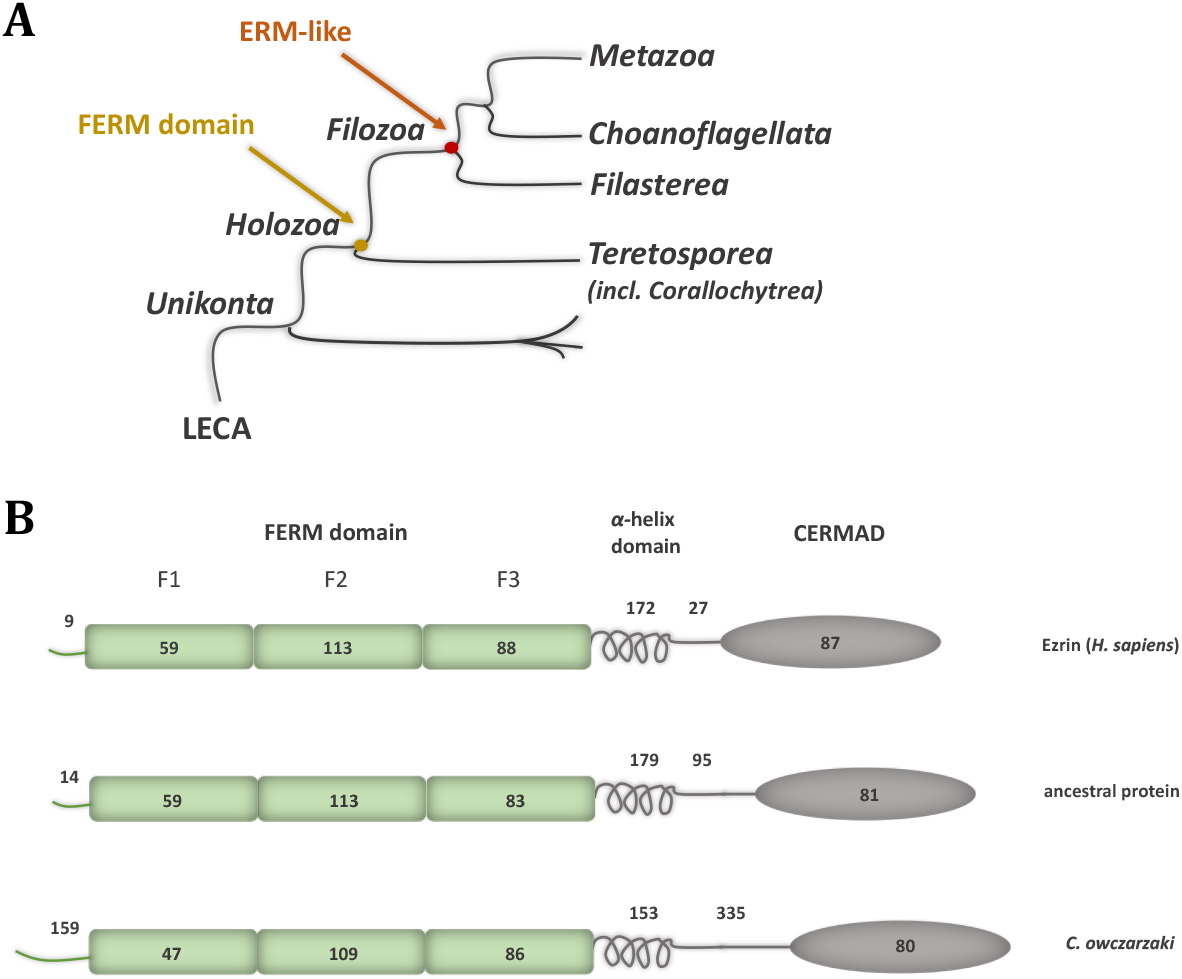
Early metazoan history of ERM+merlin protein family. A: Schematic illustration of the early ERM+merlin phylogenetic history. The arrows show first appearance of the protein structures. B: Domain structure comparison. The predicted ERM/merlin ancestral protein and the protein from *C. owczarzaki* demonstrate longer insertions between the α-helix and CERMAD. Numbers indicate number of amino acids.

We found first full-length ERM/merlin-like proteins in the closest unicellular relatives of metazoan – choanoflagellates (*S. rosetta* and *M. brevicollis),* and filasterean (*C. owczarzaki*). These proteins combine some characteristics of ERM-like (for example, C-terminal ABS) and merlin-like (multiple dispersed prolines at the C-terminal end of the alpha-helical domain, absence of the ERM-specific R_/k_EK_/r_EEL repeat within the α-helix) groups. It points out that modern merlin-like and ERM-like proteins likely emerged from the same ancestral form in the root of Metazoa. Phylogenetic reconstruction of a sequence of this ancestral form suggests that it could bind actin filaments and PIP_2_ lipid, therefore, could perform the function of mechanical linkage of the cell membrane and underlying actin filaments.

An ancestor of filasterean and choanoflagellates was likely able to form transient cell-cell or cell-surface contacts and could exploit its ERM/merlin-like protein for this purpose, as *S. rosetta* and *C. owczarzaki* probably do, as suggested by sequence analysis of their homologs. This trend could be expanded in primitive metazoans to the scaffolding function within cell-cell contacts, for example, adherens junctions in trichoplax. ERM-like protein of this early branching metazoan already possesses the key features of the ERM family: ABS and binding sites for PIP_2_, EBP50, ICAM-2, NHERF2, and intramolecular binding sites. Similarly, one study suggested that ERM proteins were involved in the development of the filopodia in metazoans (Sebé-Pedrós *et al.*, 2013). However, more sequencing data from other unicellular taxa and in vivo experiments are required to support or reject this hypothesis.

Function of the protein family in the unicellular organisms was probably limited and restricted to scaffolding, partly because of inability to activity regulation. Indeed, the auto-inhibitory interaction in the unicellular proteins is questionable, due to the presence of an extra amino acid stretch between the middle and CERMAD domains (Fig. 7B). Prediction of the tertiary structure of this insertion indicates that it is unlikely an α-helix, therefore, such inclusion could drastically change protein folding.

Further in the metazoan evolution, decreasing distance between the middle and the C-terminal domain could be one of the evolutionary modifications that facilitated ERMs’ characteristic auto-inhibitory, intramolecular binding. Therefore, our hypothesis is in accordance with the rheostat-like model of ezrin activation that ascribes the major role in this process to the alpha-helical domain (Li *et al.*, 2007). The rheostat-like manner of activation allows intermediate protein states between its inactive and active form. This multilevel manner of conformational regulation granted biochemical flexibility to ERM and merlin proteins. Consequently, they evolved more regulatory and binding sites and, therefore, acquired more functions in the cell, and eventually, became involved in signaling pathways. ERM’s intricate activity regulation mechanism became beneficial in vertebrate animals with increasing complexity of their cellular physiology and number of cell types. As a result, three ERM and two merlin paralogs diverged, acquiring some tissue specific functions in a process that can be described by the birth-and-death model (Nei and Rooney, 2005).

In agreement with the high sequence conservation among different lineages, ERM proteins also show high synteny preservation, especially within mammalian taxa, but also in such a distantly related animals as trichoplax and sponges. Surprisingly, very poor synteny conservation can be observed between ERM proteins within species. In the case of human ERMs, one syntenic genes pare for regions around moesin and radixin genes, and one – for regions around ezrin and radixin genes. Keeping in mind the mentioned conserved features of the proteins and their coding genes it is unlikely that the ERM paralogs appeared within a local duplication event. Rather the genomic regions including the ERM genes underwent significant diversification within species.

Spatial expression of paralogs is often shown to be different from the ancestral gene, that can be a sign of sub-functionalization (Glasauer and Neuhauss, 2014). Indeed, the existing RNAseq data of ezrin, radixin, and moesin demonstrate different expression patterns in different human tissues (Fig. 8). This can explain the inconsistency of some data about ezrin, moesin, radixin, and merlin roles in cancer, since the experimental results can be influenced by the cell/tissue type, momentary availability of interacting partners or ERMs’ binding sites exposure. Therefore, it is important to investigate in details the ERMs and merlin expression levels, for example, with the use of long read sequencing, and including experiments in different developmental stages. Tissue distribution of different splice isoforms is another interesting topic in ERM+merlin research: for example, in humans there are two ezrin splice variants, six – for radixin, five – for moesin, and eleven – for merlin. Unfortunately, not much is known about functions of different isoforms.

**Figure 8.**
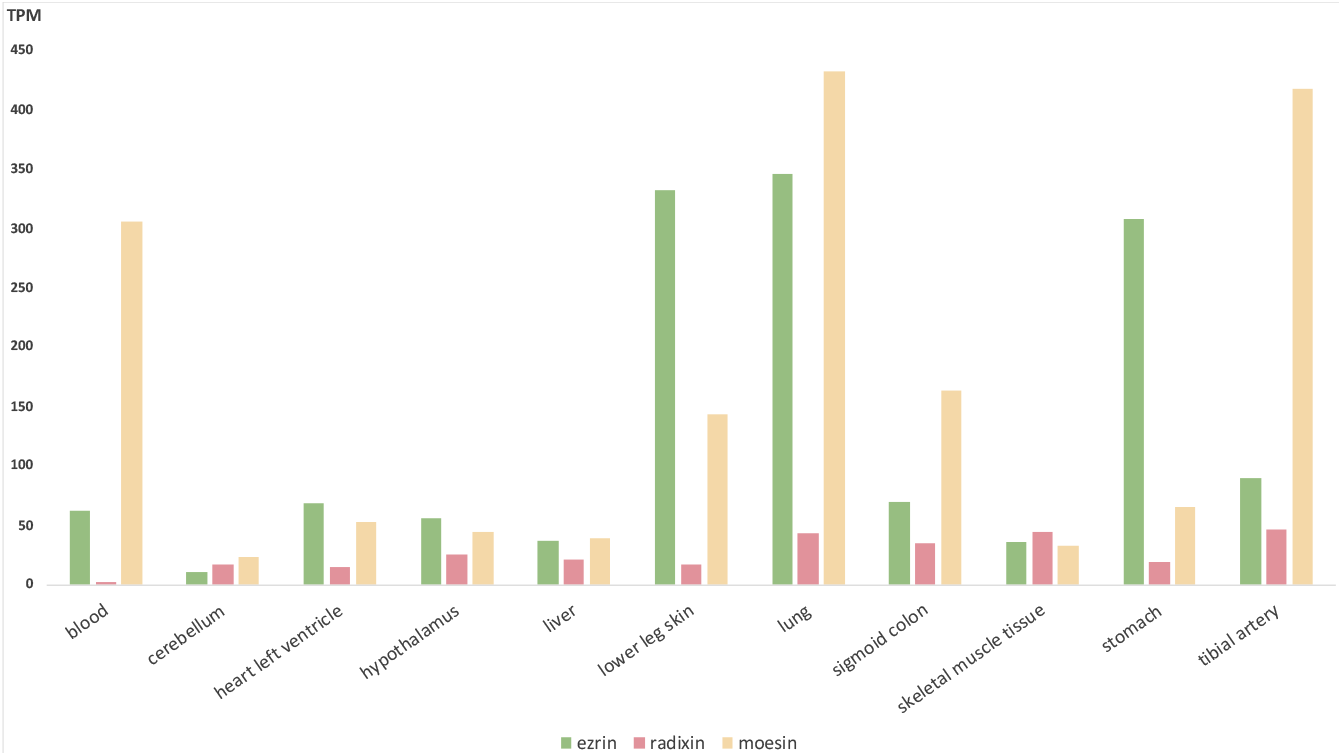
Expression levels for human ezrin, radixin, and moesin in different tissues. TPM – transcripts per kilobase million. Based on the data from Genotype tissue expression portal (GTExPortal, 2019).

Biochemical and physiological studies of the paralog specific amino acid variations are important for understanding each protein’s role in healthy and cancer cells. Based on such variations’ conservation within vertebrate lineages, we highlighted several motifs that can be candidates for such studies. For example, there is not much known about the role of the polyproline stretch in ezrin and radixin. Moesin lacks this stretch although has a structural analog (Li *et al.*, 2007), and merlin possesses multiple discontinues prolines in the homologous site. The polyproline stretch can possibly bind SH3 domain as another way of the proteins’ activity regulation (Li *et al.*, 2007) that would not be possible in moesin and merlin. Another motif of interest is located at the beginning of the α-helix and is specific to each of the proteins: EREKEQ in ezrin, EKEKEK in radixin, EKEKER in moesin, and ERTR/EKEK/EREK in merlin. This motif was earlier shown to be important for supporting the coiled-coil folding (Phang *et al.*, 2016). Next to it there is the REKEEL motif that is specific to ERMs and is absent from merlins. Another candidate is the amphipathic stretch of 14 amino acids within the α-helix region that is known to be essential for binding regulatory Rll subunit of protein kinase A (Dransfield *et al.*, 1997). This region is highly conserved in ezrin and radixin but in moesin the conservation level is only 70%. The six amino acids motif in the N-terminal end of CERMAD is H_/Q_DENxA in radixin and moesin and xxExS_/x_x in ezrin (where x is any amino acid, S/x meaning that S is conserved in half of the cases).

At the same time, ezrin, radixin, and moesin retained the set of overlapping, redundant functions, most essential for cell survival, such as organizing molecular complexes in the regions of cell-cell contacts. Ezrin is considered to be the major, indispensable paralog. Indeed, its knock out in mice causes early death of the animals. However, genomes of Tasmanian devil lack ezrin gene, at least based on the sequencing material that is available for this species at the moment. Some birds and mammals (elephant shrew) lost either radixin or moesin genes. This suggest the genetic plasticity of ERM family between different vertebrate lineages that is likely due to the conformational plasticity of the proteins.

In this work, we for the first time, to our knowledge, stress the existence of the two merlin paralogs in the vertebrate genomes: merlin1 and merlin2. Merlin1 was, apparently, lost in the Eutheria lineage, while merlin2 is present in all vertebrates. Merlin1 contains an additional amino acid stretch within its CERMAD that is absent from merlin2. Also, merlin1 has a weak sequence similarity to merlin2 in the last 30 amino acids, that is responsible for anti-tumor activity in human merlin2. Strikingly, merlin1 lacks one of the two N-terminal actin binding sites (in the F1 subdomain). These observations together with the fact that actin binding is important for merlin anti-proliferative activity (Cooper & Giancotti 2014), suggests that merlin1 is unlikely to exhibit tumor suppressive effect. It can, therefore, perform a novel, unknown function or specifically participate in cytoskeleton scaffolding, similar to merlin2 in its unfolded conformation (Lallemand, Saint-Amaux and Giovannini, 2009).

The paralog number of merlin proteins is extremely conserved among all metazoan: while ERMs’ gene number can vary as a result of lineage specific duplications, there is always only one merlin gene in invertebrates and mammals (except armadillo, who keeps, both merlin1 and merlin2) and two merlin paralogs (merlin1 and merlin2) in other vertebrates. Such conservation may signify the sensitivity to gene dosage effect for both merlin paralogs. It is unclear, though, why merlin1 was lost in the most of mammalian lineages and what protein took over its function. One could also speculate that this paralog was lost together with the organ/tissue-specific function that are present in Amphibia, Sauria, and fishes, but not in Mammalia.

At the same time merlin2 seems to be evolved specifically to counteract cellular dysregulations leading to cancer. It is likely that within evolution merlin2 paralog acquired additional to scaffolding function to contribute to evolving of the intricate anti-cancer protective system in vertebrates. Its increased in comparison to merlin1 importance is mirrored in higher synteny conservation and lower rate of nonsynonymous substitutions appearance. One can say that merin2 protein is one of the features that were evolved by the animal cell as adaptation to complex multicellularity.

Emergence of new proteins and new protein functions is an important question in evolutionary biology, and fate of paralogous forms is probably one of the least understood aspects of the process. Based on sequence comparison and phylogenetic reconstruction we hypothesized the way the ERM+merlin protein family could have gone from the first appearance of the FERM domain in holozoans to the functionally multifaceted group of five homologs with tissue specificity. We propose that the three ERM paralogs retained in the vertebrates due to their conformational plasticity that appeared to be beneficial in the conditions of the vertebrate evolution: increased complexity of organisms’ physiology and biochemistry of the cells. Merlin1 paralog is for the first time discussed here, and suggested to perform a yet unknown function specific to non-mammalian vertebrates. Merlin2 is the most interesting example of this protein family evolution, as it seems to be specifically adapted in vertebrates to anti-cancer protection.

## Materials and methods

### Data collection

The amino acid sequences of ezrin, radixin, moesin, and merlin were collected using BLASTp (Altschul *et al.*, 1997) with the human protein sequences as queries (ezrin NP_001104547.1, radixin AAA36541.1, moesin NP_002435.1, merlin NP_000259.1) against non-redundant (nr) protein sequences collection at the NCBI database. The selected sequences were manually monitored to exclude database duplicates, splice variants, truncated sequences and to reconstruct correct protein sequences when needed. If several splice variants were described for an organism, the longest one was chosen for the analysis. Only the sequences that spanned all three ERM domains were selected for the analysis. PFAM (Finn *et al.*, 2006), InterProScan (Jones *et al.*, 2014) and CDD domain search (Marchler-Bauer *et al.*, 2015) were used for domain structure analysis and verification.

Taxa selection was done based on the following requirements: 1) every described order of Metazoa should be represented by one species, although some exceptions were made in order to balance taxa representation; 2) whole genome or transcriptome of a representative species should have been sequenced and available; 3) high quality of the genome assembly annotation. In the case if no representative genome was available for an order, tBLASTn search was run against all available nucleotide sequences for that taxa. Search for possible homologs of ezrin, radixin, moesin, and merlin was performed among other Opisthokonta (Holozoa, Nuclearia, Fungi) and Amoebozoa, Excavata, Archaeplastida (includes green plants), SAR cluster (Stramenopiles, Alveolata, and Rhizaria), and other protist groups (refer to the tree of Life scheme (Adl *et al.*, 2012)). Prokaryota and Archaea nucleotide sequences were scanned for the whole length proteins or only FERM domain using tBLASTn search against nr nucleotide collection at NCBI. The taxonomic structure describing the final dataset can be viewed at the Supplementary (Table S1) and is based on the topologies employed by NCBI Taxonomy database (Federhen, 2012) and the Tree Of Life project (Letunic & Bork). The taxa variety will be further discussed in terms “vertebrate” and “invertebrate”, the later including the rest of Eumetazoa. The sequences were retrieved by July 2018; data for vertebrates were updated in May 2019.

### Reconstruction of the ERM+merlin phylogeny

Multiple sequence alignment was generated using MAFFT software (Katoh *et al.*, 2002) with the PAM70 (Dayhoff, 1965) substitution matrix, as defined by ProtTest3 (Darriba *et al.*, 2011), and manually edited to remove uninformative columns. CLUSTALX (Larkin *et al.*, 2007) and Geneious (Geneious, 2019) were used for the alignment visualization. Maximum Likelihood (ML) phylogenetic trees were build using RAxML tool (Stamatakis, 2014) with the parameters estimated by running RAxML parameter test (Stamatakis, 2015). As a result, the PROTGAMMALG model was chosen, where GAMMA model estimates the substitution rate between sites and LG is amino acid substitution matrix (Le and Gascuel, 2008). The statistical support for the tree clustering was calculated by running 1000 bootstrap replicates. The resulting trees were inspected and edited using iTOL online tree viewer (Letunic and Bork, 2019) and FigTree software (Rambaut A., 2019).

A reduced data set (Supplementary, Table S2) was used to reconstruct a tree for analyzing evolutionary relationships between the proteins from the unicellular organisms and metazoans with the same model as described earlier. Ancestral sequence reconstruction was performed for this data set using rooting option and defining marginal ancestral states option by RAxML with the model PROTGAMMALG.

MEGA (Kumar, Stecher and Tamura, 2016) software was used for an alternative tree reconstruction for neighbor-joining and parsimony algorithms with the 500 bootstrap replicates and LG amino acid substitution matrix. Bayesian inference method was also applied on MrBayes tool (Ronquist *et al.*, 2012) under LG matrix. Six chains were run for 3000000 generations, every 1000 generations trees were sampled in two runs. The first 25% of trees were discarded before constructing a consensus tree.

### Protein sequence analysis

Tertiary structure of polypeptides was predicted by PEP-FOLD3 (Lamiable *et al.*, 2016). Estimation of proteins’ biochemical and biophysical characteristics from their amino acid sequences was done with ExPASy ProtParam (Gasteiger *et al.*, 2005). Conserved amino acid motifs were analyzed throughout major metazoan lineages using MEME suite (Bailey *et al.*, 2009) on the selected set of protein sequences (see Supplementary, Table S2).

### Testing for positive selection in vertebrate ezrin and merlin

Codeml tool from PAML software (Yang, 2007) was run under model = 1 to test whether ratio of nonsynonymous-synonymous substitutions varies among tree branches. Twice difference between log likelihood of the alternative and the null (the ratio is the same for all branches) hypotheses were compared to χ^2^ distribution. Several lineages then were tested for positive selection: model = 2, NSsites = 2, fix_omega = 0, omega = 1. The null hypothesis in this case was estimated under parameters model = 2, NSsites = 2, fix_omega = 1, omega = 1. As well, χ^2^ distribution test for statistical support was used. Each codeml estimation of alternative hypothesis was run twice with varying parameter ‘omega’ to test for contingency. Parameter ‘cleandata’ was set to 0 allowing for retaining gaps in alignment.

PAL2NAL tool (Suyama, Torrents and Bork, 2006) was used to generated nucleotide alignments based on amino acid multiple sequence alignments and the corresponding mRNA sequences. The mRNA sequences were extracted from NCBI data base by using efetch command from E-utilities suit (Sayers, 2018). The phylogenetic trees for ezrin and merlin protein set from vertebrate animals were built as described earlier and can be found in Supplementary file.

### Synteny analysis

A BLASTp strategy was applied to analyze syntenic relationships between genes coding for the ERM and ERM-like proteins in different species. Sequences of the proteins encoded by the genes surrounding ezrin (seven proteins), radixin (three proteins), and moesin (four proteins) from *H. sapiens* within the region of 1 Mb were used as query in BLASTp search against all proteins in the following species: *M. musculus, D. rerio, D. melanogaster, S. pistillata* (cnidarians), *T. adhaerens,* and *A. queeslandica* with expect threshold of 1e-10. The analysis was repeated with the similarly selected proteins from *D. melanogaster* as a query. Intraspecies synteny analysis for *H. sapiens* was done in a similar manner, but with only human proteins in the target, with expected threshold of 1e-10.

For synteny analysis of the merlin gene surrounding, two species with both merlin paralogs were chosen: *D. rerio* and *G. gallus.* All proteins, which genes are located within 1Mb of merlin1 or merlin2 genes, were taken as queries. BLASTp searches with expected threshold 1e-10 were run against all proteins from the corresponding chromosomes: chromosomes 5 (has gene coding for merlin2 protein) and 21 (merlin1) for D. rerio, and chromosomes 15 (merlin2) and 19 (merlin1). Synteny Database (Catchen, Conery and Postlethwait, 2009) was used for processing the synteny data for *H. sapiens* and *D. rerio.*

Custom python, perl and bash scripts were used for data processing.

## Acknowledgments

This work was supported by the Institute of Bioinformatics, University of Muenster. The open access publishing is supported by the University of Muenster.

## Competing interests

The authors declare no competing interests.

All authors have read and approved the final version of the manuscript.

