## Supplementary material for "Emergence and evolution of the ERM proteins and merlin in metazoans": Suplemental file

Supplemetary Sequence 1.

>FERM-like domain in *C. limacisporum*

PGIIDLWTPDVKFAVRVYPELTADALLQYVCRALGLRETWYFGVNRITPTGHRVYVPVKKKMQKFLEKDR

GITVEICIACYPDTIDRLVLPETLNLFFELVSRDVMDGTLRVRPEDSIYLAAHMLHIRQGAYNTKRDKSG

FIAVKDYIPEYVLNEIALSDDQWEERIVSWYSDLKGLNETEAKKEFLHHCSVKCVNFGTFFFPVKTKKKS

AYDIALDINGLYLFDKDNRLKPLVSYLWGEIKSMSYGKTEFALKFNDRNNKEVKLIADEETLTGVLWSQA

VKHYHLWTESRQPDDLVLSQMKTLYKKNLQKREAYLQRMQWEIALRNQSEDMLR

Supplementary Sequence 2.

>predicted sequence for ERM+merlin ancestral protein form

MLRRGMPKPINVRVVTMDAEMEFSIQPKTTGKQLFDQVVRTIGLRETWFFGLQYTDTKGFVAWLKMDKKVLSHDVRRMDKSDKEEPLQFHFRAKFYPEDVSEELIQDITQHLFFLQVKEAILSEEIYCPPETCVLLASYAVQAKYGDYNPEVHKPGFLSHERLLPQRVLNQHQMTPEMWEERITAWYAEHRGMAREEAMMEYLKIAQDLEMYGVNYFPIKNKKGTPPDLWLGVDALGLNIYEKDDRLTPKIGFPWSEIRNISYNDKKFVIKPIDKKAPDFVFYSPRLRINKRILQLCMGNHELYMRRRKPDTIEVQQMKAQAREEKARKQMERQRLAREKQAREEAEREKEQMMRTRDELELRLKQCEEEAKQAEQALSRSEETAQLLAEKMKMAEEEAARQERERLLAQRAKREAEEQMTRMKKTQTRTAEEKREMLAKVRVEAEEAAQRMADEAERREQEAEELQQDLMKAKEQQVRAQHCLMNCVAASPPTVFAYPPYEPIPAPLPPDIPSFNSLIGDSLSFHVQETLQDMGVDTTQFSAELSHPTLSDAPGSSWSSSGVGAGVGVGQVGHGVSGGTEVRRGVYTAGVQAGPRDLSARPPGDGKGGPSWEVGDRDEEKRVEYMEKNKRLQQQLKSLSTDLEALRDPEKETAFDRLHNENTDRQGRDKYKTLRKIRQGNTRQRIAEFEEM

Supplementary Tree 1. The ERM+merlin phylogeny tree, Newick format with branch lengths and bootsrap:

((((Ca.lu_ezr_NP_001273918.1:0.00560144330235105587,(Eq.ca_ezr_XP_001492102.4:0.01984773245666791863,Su.sc_ezr_XP_013847913.2:0.03389229554698616059):0.01092512380428336399[51]):0.01161243610764701138[35],((Lo.af_ezrI1_XP_010597143.1:0.02325101084759792838,(El.ed_ezrI2_XP_006901976.1:0.06832630119933809365,El.ed_ezr_XP_006882827.1:0.00352449286508549299):0.02309443860683416463[99]):0.00456393073557397576[49],Rh.si_ezr_XP_019605046.1:0.01624517510924880156):0.00257413278383475386[13]):0.00676098908063847096[20],((((Al.mi_ezr_XP_014452266.1:0.06612204751959706306,(Ti.gu_ezrI1_XP_010222985.1:0.02427897813896057883,((Ga.gl_ezr_NP_990216.1:0.03248287739875620195,Ac.ch_ezrI1_XP_009073620.1:0.03551561790100313254):0.00698096175242226766[58],(Ap.fo_ezrI1_XP_009279722.1:0.00981460281500678641,Ca.an_ezrI1_XP_008495098.1:0.01466482303929032664):0.00092569124333392557[56]):0.01887238234392485392[77]):0.02628702953867440711[97]):0.02043969307232030436[74],((Th.si_ezr_XP_013913154.1:0.06768622003609583992,(Ge.ja_ezr_XP_015262280.1:0.03116006127698270345,Po.vi_ezrI1_XP_020665121.1:0.03483638498760522478):0.00825172718296463166[59]):0.02325981061308218453[93],(Ch.my_ezr_XP_007059160.1:0.02072624900168404741,Ch.pb_ezr_XP_005291597.1:0.01149633661853351196):0.03409446246882285453[100]):0.00882408197639545332[36]):0.02522719673410724409[73],(((Rh.ty_ezrREC_reconstracted:0.15887744896769076530,Ca.mi_ezrI1_XP_007892075.1:0.19389864589934024952):0.09743238469174179961[99],(((Le.oc_ezrI1_XP_006625982.2:0.06322505736946118504,((Bo.pe_ezrL_XP_020795562.1:0.21795214818704591875,((Da.re_ezr_NP_001025456.1:0.10315896834152414596,Cl.ha_ezr_XP_012676426.1:0.12783574756202498168):0.01720204136703485168[53],(Es.lu_ezrIX1_XP_010876710.1:0.06732246422677229392,(Sa.sa_ezr_XP_013999466.1:0.04191701742770672506,Sa.sa_ezrL_XP_014060138.1:0.01554200995431966222):0.01865786104994856739[97]):0.07191836971596356209[100]):0.01068196922720344305[44]):0.00913520162686813457[15],Sc.fo_ezrL_XP_018599264.1:0.09093292345204072313):0.02572149465793910350[34]):0.00992246738016824489[28],((Bo.pe_ezrL_XP_020777613.1:0.38235604790090160776,Sc.fo_ezrL_XP_018584888.1:0.14621212613256173718):0.03170523120238319392[46],(Cl.ha_ezrL_XP_012681700.1:0.25280985873486311322,(Py.na_ezrL_XP_017567344.1:0.13170844972676715168,Da.re_ezr_NP_001018326.1:0.14681037371116251178):0.03342964380854901441[54]):0.05350444156681855024[75]):0.03689636958824223795[51]):0.05975982822011949303[93],(La.ch_ezr_XP_014352395.1:0.19185549386934644400,((((((Br.be_rdxI1_XP_019624220.1:0.35598461443987455821,((Mo.br_hyp_XP_001743289.1:0.54143726948031944968,Sa.ro_ermL_XP_004994097.1:0.37636502141280159250):0.43381697382161088505[100],(((((((((Ca.mi_mrlI1_XP_007906650.1:0.03992562430471516960,Rh.ty_mrl_XP_020377697.1:0.09834182921350437256):0.04698212931149682581[100],((La.ch_mrlL_XP_014347210.1:0.14422783391773064121,(((Ch.my_mrlL_XP_007054263.1:0.01810292051777665986,Ch.pb_mrlI1_XP_008177110.1:0.00631907698767050080):0.03414267700159934887[100],(((Or.an_mrl_XP_016082473.1:0.23791137751440091797,Ph.ci_mrlI1_XP_020862733.1:0.18250593678937404585):0.04897009340992070081[66],(Ge.ja_mrlL_XP_015269113.1:0.10613917430802922992,(Th.si_mrl_XP_013923662.1:0.07384594575643031733,Po.vi_mrlL_XP_020659605.1:0.29523901358881382562):0.02911804300986619090[75]):0.12287357863696111993[100]):0.02967708650833790138[49],(Al.mi_mrlI1_XP_006260247.2:0.04395864109800150937,(Ga.gl_mrlL_XP_004946574.1:0.02878717584541263391,(Ap.fo_mrl_XP_009285644.1:0.01128111163711928953,Ti.gu_mrlL_XP_010225777.1:0.05383685682278568918):0.00678340892101360858[75]):0.04338443737357662588[100]):0.01694924260894524817[82]):0.01529664944490407609[37]):0.09018925256685925151[80],Xe.la_mrlL_XP_018102425.1:0.75472244576179536857):0.16588768390439220801[100]):0.04693067629121523004[97],(Le.oc_mrlI1_XP_015222944.1:0.07852825643975203107,(Es.lu_mrlIX1_XP_010903298.1:0.01959983255309574735,(Da.re_mrl_NP_998116.1:0.07115249878216504953,Bo.pe_mrlL_XP_020778484.1:0.07285269860959767141):0.01736689755381155137[40]):0.05762582807295340420[96]):0.10645612680062434274[100]):0.04052923200021223138[80]):0.22040134434640093475[100],((La.ch_mrlI1_XP_014351584.1:0.06921581463077901286,((Al.mi_mrlI1_XP_014449095.1:0.00713936734838541056,(((Ge.ja_mrlI1_XP_015280792.1:0.02201845679720874221,Po.vi_mrlI1_XP_020656472.1:0.01815829432365511156):0.01580630295985583147[99],(((Mo.do_mrlI1_XP_007490580.1:0.01239305299875131071,(Ph.ci_mrlI1_XP_020852055.1:0.00571548192022891706,Sa.ha_mrl_XP_012399463.1:0.04625750724277079284):0.00801027925859523239[50]):0.01347264917557965125[64],(El.ed_mrlI1_XP_006894826.1:0.00300002145768606641,(Da.no_mrlI1_XP_004461821.1:0.00687367966795787537,(Ho.sa_mrlI1_NP_000259.1:0.00229078336765698148,((Rh.si_mrlI1_XP_019598638.1:0.00458400243696887189,Lo.af_mrlI1_XP_010597054.1:0.00685318959620390133):0.00000100000050002909[13],((Eq.ca_mrl1_XP_001498910.1:0.00453656688984304529,Ca.lu_mrl1_XP_534729.2:0.00000100000050002909):0.00227428589757489030[65],(Ra.no_mrl_NP_037325.1:0.03759479768391412596,Su.sc_mrl1_XP_003483494.1:0.00000100000050002909):0.00227959897550708730[36]):0.00000100000050002909[6]):0.00000100000050002909[3]):0.00000100000050002909[43]):0.00382371885758110057[55]):0.03385466457784968436[70]):0.01056531314465571948[91],(Xe.la_mrl_XP_018115976.1:0.01294500860876077533,Xe.la_mrlSH_XP_018093062.1:0.01661566619253309479):0.10955156792552753209[100]):0.00821802914742108051[56]):0.00468656235878304236[27],(Ch.pb_mrlI1_XP_005278881.1:0.00000100000050002909,Ch.my_mrl_XP_007068562.1:0.06683950187023904310):0.00893193796630869195[96]):0.00501945050127198394[39]):0.00000100000050002909[16],((Ga.gl_mrl_NP_989828.2:0.00000100000050002909,Ca.an_mrl_XP_008499723.1:0.00242329329875604288):0.00219019146322837251[96],Ac.ch_mrlI1_XP_009070304.1:0.02628069329665234238):0.01092895463338867078[97]):0.06571408741008548382[99]):0.02868493690808148616[90],((Le.oc_mrl_XP_006640249.1:0.01711502411416928118,((Py.na_mrl_XP_017560541.1:0.05175715188680301421,(Sc.fo_mrl1_XP_018596058.1:0.11143656143407727754,Da.re_mrl_NP_001122179.1:0.08626277999943927910):0.01952819580409252687[48]):0.01067170439118923092[40],(Cl.ha_mrlIX_XP_012671441.1:0.08374540991227887032,(Bo.pe_mrl_XP_020774724.1:0.15083021920628256196,(Sa.sa_mrl_XP_014026700.1:0.03935119978490241033,Es.lu_mrlIX1_XP_012988213.1:0.05057894361314441839):0.05520542926172854886[100]):0.03145701170269069036[95]):0.00762911502251461919[50]):0.04801665922071329806[96]):0.05422506898191362112[100],Ca.mi_mrl_XP_007900504.1:0.17443085351428211371):0.01777546904068980430[74]):0.11171671990361067839[100]):0.27696145132297428360[100],(Hy.vu_mrlL_XP_012562032.1:1.34276478097717166804,Mo.te_mrl_BAF49216.1:1.01231893713392073764):0.23465177504283024623[53]):0.05818096108104554853[28],(((Ac.pl_mrlL_XP_022093213.1:0.20515825621764019471,St.pu_mrl_XP_011683075.1:0.33834094768360889471):0.22892430831689147830[100],Sc.ko_mrl2_NP_001164711.1:0.34941122313976918923):0.04670110804068505067[30],(((Li.an_mrlL_XP_013419248.1:0.34665059673243414640,((Lo.gi_hyp_XP_009051406.1:0.26491524267272614779,Bi.gl_mrlL_XP_013091480.1:0.36348131762221130847):0.07032159685788487435[95],(Cr.gi_mrlI1_XP_011437863.1:0.18307701435634687881,Mi.ye_mrlL_XP_021374387.1:0.23335730464222240177):0.10123686551959719393[100]):0.08143843733238992355[93]):0.06435525436933131616[85],(Pr.ca_mrlL_XP_014661700.1:0.47705746323167141920,(((As.su_mrl_ERG80906.1:1.04879637993112950767,(Tr.ps_mrl_KRY01563.1:0.02143450261936851245,Tr.ps_mrl_KRY91900.1:0.00100501568358580444):0.54560230539088150348[100]):0.29730944652680962870[96],(Ra.va_hyp_GAU96524.1:0.28324073256307419344,Hy.du_mrl_OQV19900.1:0.49333576146967916820):0.91192088267376991695[100]):0.09208813219744572953[22],(((Tr.me_mrlL_OQR68523.1:0.79014481661135438362,Te.ur_mrlL_XP_015784229.1:0.38687967063495709574):0.02716561213446477135[12],(St.mi_mrl_KFM73866.1:0.03105552517312464705,Pa.te_mrlL_XP_015909486.1:0.09155168021878073992):0.17898159338686092656[100]):0.08228541757390848976[38],((Or.ci_mrl_ODN01088.1:0.41700726400704951624,(Da.pu_hyp_EFX76345.1:0.35073374828149972426,(Gr.bi_mrl_BAN09208.1:0.12584062747102381374,((Ci.le_mrl_XP_014247697.1:0.17451611936539543346,(Dr.me_mrlIA_NP_523413.1:0.91871950944653668625,Sp.li_mrlL_XP_022831808.1:0.23595054169804055566):0.06529208535257047252[41]):0.02598547488324036772[25],((Ni.ve_mrl_XP_017787144.1:0.24206222781157010759,Pe.hu_mrl_XP_002427487.1:0.24298372898945835852):0.02635277575065344755[25],(Zo.ne_mrl_XP_021921812.1:0.06279664737157988896,Ca.fl_mrl1_XP_011258088.1:0.10201484408944168358):0.01968440680241427543[45]):0.02142523964021291610[28]):0.02849000329434924275[26]):0.14113479054608599195[71]):0.02693899599344085072[20]):0.04738225662093796531[24],Eu.af_mrlL2_XP_023328819.1:0.52966827420061068921):0.06625779819373470159[46]):0.04758493814814217238[33]):0.06391130535217076636[57]):0.06235095027491030506[57]):0.11573682208285336614[62],(Br.be_mrlL_XP_019621260.1:0.30860392172976663927,St.pi_mrlL_XP_022798867.1:0.74782012720984614162):0.06382979027583057796[25]):0.03953043072471019298[20]):0.05422358120998738151[14]):0.11956879160011087138[59],Am.qu_mrlL_XP_011410150.1:1.35325659943545084651):0.13393961021879660644[71],(Sa.ro_moe_XP_004991962.1:1.18461811522163995569,Ca.ow_rdx_XP_004364665.2:1.54573586188841227695):0.15869001273510177641[63]):0.27292618137734497852[88],In.li_hyp_OAF67650.1:2.97049676232722958957):0.22080411311529973828[28],((St.pu_moe_XP_011674768.1:0.24116351703925903438,Ly.va_moe_P52962.1:0.33935582870775193864):0.46138941786877590845[100],(Ex.pa_ermH_XP_020900768.1:1.03182434931372402076,(Sa.ro_cytsk_XP_004997754.1:0.46945736982181451857,Mo.br_hyp_XP_001746613.1:0.73145109953294951133):1.40222268586173770544[100]):0.18735666920113269729[12]):0.04795889913135345517[5]):0.05674184792858815579[2]):0.04891966200444984592[2]):0.09022370940055894628[2],Sc.ko_rdx_XP_002733683.1:0.37151171813234157293):0.04801688104597026663[1],(((St.pi_moeL_XP_022781886.1:0.73549062254598795985,(Ci.in_ermL_NP_001027599.1:0.50466829196199936014,Mo.te_erm_p_BAF49215.1:0.59636058324651841644):0.12950327047337031883[86]):0.08193395126715030674[22],Ac.pl_rdxL_XP_022107608.1:0.55146054996240867485):0.09562014370394752993[11],((Li.an_ermL_XP_013416507.1:0.34179473741420302035,Li.an_rdxL1_XP_013416553.1:0.26579638181961906529):0.16026327273409909924[97],(((((Ma.li_hyp_PAA60758.1:0.77942458967896510735,(Cl.si_rdx_GAA55417.1:1.78558528762960921910,Ec.gr_erm_EUB64035.1:1.41637764738166582745):0.10581109648015807334[19]):0.05410361278030542675[14],(Ex.pa_rdxL_XP_020900767.1:0.90163800123917892115,Am.qu_rdxL_XP_003386140.1:0.45913890169150245457):0.11631277144773304044[19]):0.08374142178881337217[2],(((((((Pe.hu_erm_XP_002429229.1:0.23239516690364842022,Zo.ne_ermH_XP_021936488.1:0.07378279332516306244):0.01358434746036349915[32],(Ca.fl_ermL3_XP_011257675.1:0.17897548450379471840,Ci.le_ermI1_XP_014262198.1:0.13979846084041405718):0.02830269451439463776[60]):0.03130220837195733796[67],(Sp.li_ermL1_XP_022825687.1:0.18675030416015148127,(Ni.ve_ermI1_XP_017779440.1:0.16506693050773588172,Dr.me_moeIB_NP_727290.1:0.15630215225954705027):0.02792410709320587367[75]):0.02077133357361101926[68]):0.15618381387560353879[93],(Da.pu_hyp_EFX71611.1:0.18606686064494307176,Or.ci_erm1_ODN03767.1:0.18717122039451322690):0.04539821392346667789[61]):0.01884519104290724098[28],(Hy.az_ermI1_XP_018006433.1:0.20735007621203882522,((Ni.ve_ermI1_XP_017779445.1:1.59556203927790773989,Eu.af_ermL1_XP_023343721.1:1.03097173232983130298):0.23960445130044377704[28],Eu.af_ermL1_XP_023321842.1:0.28726886047211730446):0.01204174310584399561[3]):0.04259896667937877052[4]):0.08315338846073058732[23],(Tr.me_erm1L_OQR70766.1:0.43968641835504690407,((Sa.sc_erm1L_KPM03270.1:0.32914527432258017248,Te.ur_LermH_XP_015785745.1:0.23525319117379397960):0.04811227682526531685[44],(Pa.te_ermH_XP_015929578.1:0.15741553229113672274,(Li.po_ermH_XP_013785124.1:0.02414330633503678977,(Li.po_ermH_XP_013781090.1:0.08490152276658333164,Li.po_ermH_XP_013794693.1:0.08806980690200635897):0.02328668364671377874[49]):0.11316801696366250718[99]):0.04111932756926470200[75]):0.03299970341782153260[31]):0.04068099818764757403[64]):0.07210653145904805106[26],((Hy.du_ermL1_OQV15172.1:0.27098439103693700014,Ra.va_hyp_GAU99524.1:0.16174068225729829051):0.32517312213066362769[100],((Ca.el_erm_NP_491559.1:0.39318434039142291514,Lo.lo_moe_XP_020304013.1:0.20496471052874606911):0.21152799473583486223[100],Tr.ps_ermL1_KRY90359.1:0.39014018485643131573):0.21174976692377331378[99]):0.06505975771086830450[60]):0.18342789001627513024[53]):0.06168587099598625556[3],(Cr.gi_rdxI3_XP_011453118.1:0.34887128147955992485,(Mi.ye_rdxL_XP_021360349.1:0.71234108063404644184,Bi.gl_rdxL_NP_001298197.1:0.45161306966721714851):0.07138760499473781329[23]):0.11014246564337672185[27]):0.04661630808419102434[1],(Oc.bi_moeL_XP_014769368.1:0.62259006730422949971,(((He.ro_hyp_XP_009017370.1:1.37023376972196131440,He.ro_hyp_XP_009010884.1:0.25961068521757796335):0.12182375710581586081[63],He.ro_hyp_XP_009021945.1:0.39301629657828596187):0.12127969350509064383[88],Pr.ca_ermH_XP_014672976.1:0.64717730303336851172):0.04209659650738673681[40]):0.03477889935557043621[12]):0.04806144976728433937[3]):0.08626538700144489868[5]):0.05192742095680791953[1]):0.06282779027571164243[7],Tr.ad_hyp_XP_002112848.1:0.83389040892784638270):0.06411098514883496746[46],Hy.vu_rdxL_XP_002160112.1:0.53141198280356638506):0.25368803486435331784[97],(((((((Ch.pb_moe_XP_005279574.2:0.00818473603705250838,Ch.my_moe_XP_007060247.1:0.00394764173647085392):0.01224973440660916124[96],(((Th.si_moe_XP_013928247.1:0.04607702553353398151,Po.vi_moeI1_XP_020640009.1:0.02422283657054280157):0.01602116382276986759[90],Ge.ja_moe_XP_015277108.1:0.04930539316845845149):0.01390638642267249050[62],(((Or.an_moe_XP_007659466.1:0.13660891209930711709,((Ra.no_moe_NP_110490.1:0.01858188286796397248,(Da.no_moe_XP_004470439.1:0.01266679425815336359,(Ho.sa_moe_NP_002435.1:0.00241512899876632908,((Su.sc_moe1_XP_013841673.1:0.01483977980513038163,Ca.lu_moe2_XP_848336.1:0.01213434523804400242):0.00000100000050002909[14],(Eq.ca_moe_XP_001504911.1:0.00483750523312235364,(Lo.af_moeI2COR_XP_010591044.1:0.01208594274361066839,Rh.si_moe_XP_019573069.1:0.00967995979928567944):0.00000100000050002909[15]):0.00241566445383809307[41]):0.00245510129378259234[36]):0.00222435121676362512[28]):0.00893057675029782644[91]):0.01866766865658819538[94],(Mo.do_moe_XP_007507017.1:0.09520007876892938592,Ph.ci_moe_XP_020822867.1:0.00000100000050002909):0.02516974809468296853[99]):0.01319232399994966706[71]):0.01582177940200931582[75],(Ap.fo_moe_XP_009286260.1:0.00000100000050002909,(Ca.an_moeI1_XP_008499411.1:0.00688400413275106794,Ti.gu_Lmoe_XP_010211249.1:0.00480496049317457111):0.00286103120799135156[27]):0.03304934998993157985[98]):0.01300666853457765545[44],Al.mi_moe_XP_006270176.1:0.03777926540030774466):0.00185273922250953548[11]):0.01246950753146941482[27]):0.06009597907245010223[99],Xe.la_moeI1_XP_018088399.1:0.11646467616163175274):0.03819023460316853941[99],La.ch_moe_XP_014345259.1:0.15573458655087341063):0.03752436575918540601[99],(Le.oc_moeI1_XP_006632824.2:0.01986011015840139990,((((Cl.ha_moeL_XP_012689836.1:0.11417973709707739116,(Da.re_rdx_NP_001243180.1:0.09793305221717317488,Py.na_moeL_XP_017568853.1:0.06268389429461418416):0.02340701079976423987[92]):0.02557342899821896456[94],(Bo.pe_rdxL_XP_020778213.1:0.13914278890910117270,(Es.lu_moeIX1_XP_010892630.1:0.06969897801206585697,Sa.sa_moeL_XP_014054227.1:0.04417812217566441380):0.05747182092337423803[99]):0.05466662233478720989[99]):0.13770698044016260742[100],Sc.fo_moeL1_XP_018592896.1:0.15960396028855380890):0.02209258325364696698[87],(((Bo.pe_moe_XP_020789144.1:0.08464888958973178223,(Cl.ha_moe_XP_012678436.1:0.07774043274969841266,(Sa.sa_moe1_XP_013993006.1:0.04095833273674282654,Es.lu_moe_XP_010863311.1:0.01070287517234460098):0.04008836031347848966[99]):0.01162170455813428364[60]):0.01881752188024586950[64],(Py.na_moeL_XP_017538287.1:0.02750587215134049288,Da.re_moe_NP_001004296.1:0.04959597452040392929):0.01105158037410333520[61]):0.00324675705265897859[30],Sc.fo_moe1_XP_018584046.1:0.05670313408743140465):0.02625746042677143746[97]):0.01333519816645204956[76]):0.06310256481408756113[100]):0.05125167774470447413[99],(Rh.ty_rdxL_XP_020386890.1:0.18373653683903598544,Ca.mi_moeI1_XP_007891635.1:0.13165857987864978962):0.05917043252275761001[100]):0.10573120511725418724[100],((Ca.mi_rdxI1_XP_007903800.1:0.04158666842966731586,Rh.ty_rdxL_XP_020367415.1:0.05643005180332536647):0.03845021104138768298[86],((((Le.oc_rdx_XP_015219579.1:0.19774807106995032080,(Sc.fo_rdx1_XP_018595413.1:0.03083043496306653655,((Py.na_rdx_XP_017565517.1:0.05933129237758771185,Cl.ha_rdx_XP_012696136.1:0.04202021115414941721):0.01807049779762404801[46],((Bo.pe_rdx_XP_020793812.1:0.07989519356317893728,((Sa.sa_rdxL1_XP_014007263.1:0.00724279243717377511,Sa.sa_rdxL1_XP_014005960.1:0.01250700831971300955):0.03032277044201235616[100],Es.lu_rdx_XP_010884704.1:0.01829294790754958100):0.04054241989484316105[98]):0.04074407482055485835[80],(Bo.pe_rdxL_XP_020776109.1:0.29781148785327637984,Sa.sa_rdx1_XP_014020302.1:0.05741278832677987332):0.02023826546465787996[38]):0.00638323858396160765[25]):0.02110562659646542799[29]):0.01231585911667544553[25]):0.03512814697886904036[56],(Xe.la_rdx_XP_018105824.1:0.02832782479967819664,Xe.la_rdxL_XP_018101057.1:0.03771826206604965426):0.01335642765314526414[87]):0.02095985924554540081[18],(((Th.si_rdxI1_XP_013915432.1:0.02645745909806784812,(Po.vi_rdxI1_XP_020660843.1:0.02151438779106178839,Ge.ja_rdxI1_XP_015284977.1:0.01127211539643002865):0.01191644941036980494[50]):0.01371941226447214758[56],(((Ph.ci_rdx_XP_020851763.1:0.00000100000050002909,Mo.do_rdxI1_XP_016289356.1:0.02601829681968333141):0.01267794597682128308[89],((Ti.gu_rdxI1_XP_010221655.1:0.00733250720832966454,Ga.gl_rdx_NP_990082.1:0.00242655634807940066):0.00000100000050002909[17],(Ac.ch_rdx_KFP73328.1:0.01519797612811569706,(Ca.an_rdx_XP_008500566.1:0.00000100000050002909,Ap.fo_rdx_XP_009280888.1:0.00000100000050002909):0.00000100000050002909[73]):0.00242546671476557207[65]):0.00238635339704139403[63]):0.00425259833450375262[38],((Ch.my_rdx_XP_007059750.1:0.00240888207429777691,Ch.pb_rdx_XP_008167539.1:0.00000100000050002909):0.00482862924481283675[91],Al.mi_rdxI1_XP_014456169.1:0.01729977256648672704):0.00239527484311393302[41]):0.00387390906255965324[10]):0.00998456061275849842[21],(Lo.af_rdxI1_XP_010593746.1:0.00999264516622623555,(Ho.sa_rdx_AAA36541.1:0.00488794591093482354,(((Rh.si_rdxI1_XP_019570833.1:0.02964735483065432040,Da.no_rdx_XP_004481171.1:0.00221141953082016609):0.00229013071698418126[38],(Eq.ca_rdx1_XP_003365379.1:0.00000100000050002909,(Su.sc_rdx_NP_001009576.1:0.00489865736665241226,Ca.lu_rdx1_XP_022274045.1:0.00303994009436992386):0.00296487175823338320[55]):0.00298801104006937452[38]):0.00000100000050002909[10],Ra.no_rdx_NP_001005889.2:0.02971681355168074093):0.00000100000050002909[8]):0.00819552842404049076[33]):0.01295811575280413422[57]):0.00547923895605744327[28]):0.00779510588530702743[23],La.ch_rdx_XP_014350863.1:0.05116688280916430187):0.01644011167297191461[29]):0.05973219314884901932[72]):0.05662385493682602078[50]):0.16060891492971557382[100]):0.02835792198805425074[50]):0.01638842331952003462[63]):0.07426744432940977914[100],Xe.la_ezrLH_NP_001087392.1:0.20339869133877502838):0.07780810993635162154[100]):0.03470590609110564551[97],((Sa.ha_ezr_XP_003769620.1:0.03374282395184580868,Mo.do_ezrI1_XP_001372743.1:0.00578175862805154882):0.00272208501065583338[41],Ph.ci_Lezr_XP_020842368.1:0.01380435495174263914):0.01705459389735348302[89]):0.01998908309773695866[88]):0.00879071052123375427[24],(Ho.sa_ezr_NP_001104547.1:0.02231029996520883865,Ra.no_ezr_NP_062230.1:0.03215733059106781061):0.01027103796154712340[54],Da.no_ezr_XP_004462963.1:0.05290325366554535919);

Supplementary Tree 2. ML tree illustrating the early ERM+merlin evolution, Newick format with branch lengths and bootstrap:

(Choanoflagellatea1_ermL:0.27912655910410960614,(Eutheria_ezr:0.40791717949211842020,(((Eutheria_mrlI1:0.43505914662388855962,Choanoflagellatea1_moe:0.75915103767809610780):0.06661971040473635419[60],Filasterea_rdx:1.04917937076769773874):0.14109513997400438545[88],(Choanoflagellatea2_hyp2:0.47188356357813210362,Choanoflagellatea1_cytsk:0.24627930065746514443):1.00220236071595536487[100]):0.12503270961915494142[65]):0.11522322196398884775[89],Choanoflagellatea2_hyp1:0.32434340744908751741);

Supplementary Tree 3. ML tree for vertebrate ezrins, used in PAML analysis, with branch lengths and bootstraps:

((Ph.ci_Lezr_XP_020842368.1:0.00807425360331540283,(Mo.do_ezrI1_XP_001372743.1:0.00744864123133891828,Sa.ha_ezr_XP_003769620.1:0.02451521614668837612):0.00208505222941104138[46]):0.00648495219792183993[79],(((((Ch.pb_ezr_XP_005291597.1:0.00928064866297956606,Ch.my_ezr_XP_007059160.1:0.01572330031951145110):0.02556783744714864054[100],(Th.si_ezr_XP_013913154.1:0.05272772394800537277,(Po.vi_ezrI1_XP_020665121.1:0.02781849684684373908,Ge.ja_ezr_XP_015262280.1:0.02498807710043972341):0.00570943789555732658[69]):0.02046750665206707384[97]):0.00829476389248711386[53],(Al.mi_ezr_XP_014452266.1:0.05544730292896746238,(Ti.gu_ezrI1_XP_010222985.1:0.01795146653074274762,((Ga.gl_ezr_NP_990216.1:0.02543072203240308651,Ac.ch_ezrI1_XP_009073620.1:0.02794169622261391539):0.00514058885273716796[53],(Ap.fo_ezrI1_XP_009279722.1:0.00763381701034998843,Ca.an_ezrI1_XP_008495098.1:0.01134279284416132676):0.00083494791001704962[61]):0.01497977258611864378[83]):0.01807584711686882617[97]):0.01338566707525845785[66]):0.01847258148289673829[68],(Xe.la_ezrLH_NP_001087392.1:0.15060747726232626142,((((((Es.lu_ezrL_XP_028981409.1:0.22774139095835896351,Bo.pe_ezrL_XP_020777613.1:0.26449361035623164540):0.07485925470769709544[95],((Py.na_ezrL_XP_017567344.1:0.10535314754343670651,Da.re_ezr_NP_001018326.1:0.11851753088082417342):0.03155697025841060210[71],Cl.ha_ezrL_XP_012681700.1:0.22962682065372633233):0.02645630927543668465[77]):0.01511266565740862489[77],Sc.fo_ezrL_XP_018584888.1:0.14285872785878200864):0.01199440757846485621[75],Bo.pe_ezrL_XP_020795562.1:0.17441928354161481685):0.01653051743303209434[58],((Le.oc_ezrI1_XP_006625982.2:0.05188410692447134598,Sc.fo_ezrL_XP_018599264.1:0.06349337717687149329):0.01390038510821289819[50],(Cl.ha_ezr_XP_012676426.1:0.10842821190318274738,(((Sa.sa_ezrL_XP_014060138.1:0.01232163006760782914,Sa.sa_ezr_XP_013999466.1:0.03261953778394610648):0.01241465595762415892[90],Es.lu_ezrIX1_XP_010876710.1:0.05580283138536127480):0.05746700182073909147[100],Da.re_ezr_NP_001025456.1:0.08443503849340104617):0.00897285505741565563[42]):0.01566359970013269862[63]):0.01249239153815551347[31]):0.07027281585492088645[99],(La.ch_ezr_XP_014352395.1:0.14419882884789506083,Ca.mi_ezrI1_XP_007892075.1:0.21205506260536141627):0.01465926500383295771[64]):0.05993968137061567170[100]):0.07577184326633953915[100]):0.02670913626926851164[96],((((Su.sc_ezr_XP_013847913.2:0.02618858010143197690,Eq.ca_ezr_XP_001492102.4:0.01528284533099706285):0.00788435788206337008[41],Ca.lu_ezr_NP_001273918.1:0.00461122901069683198):0.00832916345875283140[30],((Lo.af_ezrI1_XP_010597143.1:0.01502281835229918663,(El.ed_ezrI2_XP_006901976.1:0.05349915510234238419,El.ed_ezr_XP_006882827.1:0.00204951879974538381):0.01805029110339222490[99]):0.00208175327373273308[56],Da.no_ezr_XP_004462963.1:0.04236346722258913650):0.00532522675895442422[32]):0.00226596969968256758[21],(Rh.si_ezr_XP_019605046.1:0.01143219973331763284,(Ho.sa_ezr_NP_001104547.1:0.01811245434846534691,Ra.no_ezr_NP_062230.1:0.02341481372869059011):0.01265540268419285130[46]):0.00344370750559593129[22]):0.01795195692746761815[81]):0.00890472441824967723[49],Or.an_ezr_XP_028908150.1:0.05152041395556668929);

Supplementary Tree 4. ML tree for vertebrtae merlins, used in PAML analysis, with branch lengths and bootstrap:

(Ho.sa_mrlI1_NP_000259.1:0.00000100000050002909,(((Su.sc_mrl1_XP_003483494.1:0.00244729357014871482,Da.no_mrlI1_XP_004461821.1:0.00738664363861629876):0.00000100000050002909[7],(Rh.si_mrlI1_XP_019598638.1:0.00491638396976679885,(Eq.ca_mrl1_XP_001498910.1:0.00485345329444812450,Ca.lu_mrl1_XP_534729.2:0.00000100000050002909):0.00243558683121475371[68]):0.00000100000050002909[11]):0.00000100000050002909[6],(((((((Al.mi_mrlI1_XP_014449095.1:0.00726834696129703608,((Ga.gl_mrl_NP_989828.2:0.00000100000050002909,((Ap.fo_mrl_XP_019326828.1:0.03108185940959790755,Ti.gu_mrl_XP_010216370.1:0.00557829901268469368):0.00000100000050002909[52],Ca.an_mrl_XP_008499723.1:0.00260396621427923377):0.00000100000050002909[50]):0.00235704557869804293[58],Ac.ch_mrlI1_XP_009070304.1:0.03177157152496361270):0.01236554120589733116[95]):0.00250669232176025519[51],(((((La.ch_mrlL_XP_014347210.1:0.19314187132416205106,(((Mo.do_mrl_UniProtID_F7G4H7:0.35507907758700218981,(Ph.ci_mrlI1_XP_020862733.1:0.08740860398805413989,Sa.ha_mrlL_XP_023359437.1:0.04608981959422241842):0.02031469812361758748[100]):0.10417883687741331944[100],(Da.no_mrll_XP_004460756.1:0.55736041205505226781,(Or.an_mrl_XP_028938201.1:0.21658737251056672801,Xe.la_mrlL_XP_018102425.1:1.03142610158436975887):0.05851687983185931835[38]):0.01295478906568882545[21]):0.04396240544596704408[57],((Ge.ja_mrlL_XP_015269113.1:0.12165287706845229199,(Th.si_mrl_XP_013923662.1:0.09879437325867911068,Po.vi_mrlL_XP_020659605.1:0.34420723705892131150):0.03446187720352445610[70]):0.15195233440019143001[100],((Ch.my_mrlL_XP_007054263.1:0.02001840089136320069,Ch.pb_mrlI1_XP_008177110.1:0.00640788832092629556):0.05485598469823086548[100],(Al.mi_mrlI1_XP_006260247.2:0.04598539616774730693,((Ga.gl_mrlL_XP_004946574.1:0.00000100000050002909,Ga.gl_mrl_XP_004946574.1:0.00000100000050002909):0.03178589996144211283[100],(Ti.gu_mrlL_XP_010225777.1:0.06129860926443967545,Ap.fo_mrl_XP_009285644.1:0.01064848885721431798):0.00716203944850166748[94]):0.04907177921552069444[98]):0.01609200902306867284[92]):0.03114478241524107696[75]):0.02884231177262953569[54]):0.30870014819970087450[100]):0.05653373312006441942[99],(((Cl.ha_mrlL_XP_012671951.1:0.08597721406058732307,Bo.pe_mrlL_XP_020778484.1:0.06769691467657307227):0.02276412848538960851[53],((Sa.sa_mrl_XP_013985183.1:0.03159507697339678650,Es.lu_mrlIX1_XP_010903298.1:0.02265835835889883076):0.01810943026290093641[79],(Py.na_mrlL_XP_017554102.1:0.04092802510984770714,Da.re_mrl_NP_998116.1:0.06532399829111189271):0.04221142857267100312[91]):0.01355506905083203595[25]):0.02021947923000510031[67],(Le.oc_mrlI1_XP_015222944.1:0.12782151780578884792,Sc.fo_mrlL_XP_018586205.1:0.07231947299089802605):0.01978282506611444766[30]):0.13615792665358245062[100]):0.05849275069529789872[99],(Ca.mi_mrlI1_XP_007906650.1:0.04800907418425597734,Rh.ty_mrl_XP_020377697.1:0.11755839637966919442):0.05240428777264895255[92]):0.44067164763904576041[100],(((((Py.na_mrl_XP_017560541.1:0.04781985244963920384,Da.re_mrl_NP_001122179.1:0.10037315480500384746):0.01764700087369339143[38],(((Sa.sa_mrl_XP_014026700.1:0.04393091613897727454,Es.lu_mrlIX1_XP_012988213.1:0.05376528967483664573):0.05996117041825210769[100],Bo.pe_mrl_XP_020774724.1:0.16869992287775478768):0.03511730046425998869[93],Cl.ha_mrlIX_XP_012671441.1:0.09171779075332689435):0.00738637405494778007[33]):0.02984604770718326389[68],Sc.fo_mrl1_XP_018596058.1:0.11900590888213678775):0.04020406255364925713[99],Le.oc_mrl_XP_006640249.1:0.00748272598393164273):0.05785851012012827727[99],Ca.mi_mrl_XP_007900504.1:0.19987071644302628615):0.01486102532503053714[59]):0.03074929768028106594[94],La.ch_mrlI1_XP_014351584.1:0.08123962807076791359):0.06252389021231410937[99]):0.00248477954750221837[35],((Ch.pb_mrlI1_XP_005278881.1:0.00000100000050002909,Ch.my_mrl_XP_007068562.1:0.07369814919050043933):0.00999407288475155989[97],(Po.vi_mrlI1_XP_020656472.1:0.01911006898544479263,Ge.ja_mrlI1_XP_015280792.1:0.02395560161364927720):0.01773943280204617889[98]):0.00451114109487036306[43]):0.00652713077487947387[57],(Xe.la_mrlSH_XP_018093062.1:0.01586031001411454880,Xe.la_mrl_XP_018115976.1:0.01626579581860948878):0.12797312989599851640[100]):0.00596295435329354580[69],((Mo.do_mrlI1_XP_007490580.1:0.01205038258252420780,(Ph.ci_mrlI1_XP_020852055.1:0.00731571521122090576,Sa.ha_mrl_XP_012399463.1:0.05258589195058212262):0.00756536237005694945[68]):0.01510655660757304064[80],Or.an_mrl_XP_028904749.1:0.02418900119069150589):0.00922666964934595012[66]):0.03844749944613848336[94],El.ed_mrlI1_XP_006894826.1:0.00236301245850235831):0.00496186708708381673[77],Lo.af_mrlI1_XP_010597054.1:0.00735590002452357356):0.00000100000050002909[52]):0.00244307735159380842[47],Ra.no_mrl_NP_037325.1:0.04371246054736231679);
